# Molecular adaptations of activated T-cells in an inflammation-associated schizophrenia sub-group

**DOI:** 10.64898/2026.05.22.727194

**Authors:** Deepak Salem, Rowan K. O’Hara-Payne, Sarah M. Clark, Marcia Cortes-Gutierrez, Nevil J. Singh, Daniel J.O. Roche, Deanna L. Kelly, Seth A. Ament

## Abstract

Thirty-five percent of people with schizophrenia-related disorders (SRD) form a high-inflammation subgroup defined by elevated anti-gliadin antibodies (AGA+) and inflammatory proteins and associated with an increased severity of negative symptoms. However, the immune mechanisms mediating these effects remain poorly defined. Here, we characterized transcriptional signatures of peripheral immune cells in AGA+ SRD (n=7) compared to AGA-negative (AGA-) SRD (n=3) and healthy controls (HC; n=5), using single-cell RNA-sequencing (scRNA-seq) of peripheral blood mononuclear cells (PBMCs). AGA+ SRD was associated with increased abundance of T-helper-17 cells (Th17), T-follicular helper-1 (Tfh1), CD5+ B cells, plasmacytoid dendritic cells (pDCs), and several CD8+ T cell subsets, including memory and Natural Killer-T-like activated subsets. In parallel, AGA-SRD exhibited a higher abundance of several monocyte subsets compared to either AGA+ SRD or HC. Pathway analysis revealed upregulation in AGA+ SRD of JAK/STAT, type I Interferon, and IL-6 signaling pathways in distinct subset of activated T-cells. Collectively, these results define a unique T cell predominant inflammatory signature in AGA+ SRD, as well as potential targets for therapeutic intervention.

## Introduction

The inability to treat negative symptoms in schizophrenia and related disorders (SRD) (1) is the major factor driving SRD’s impact on disability and life lost. The 20 million people living with SRD worldwide lose 28.5 years of life on average, making SRD one of the top 15 causes of disability (2, 3). Experiential negative symptoms, including profound deficits in anhedonia and motivation, are particularly disabling (5, 6), in part because there are no FDA approved treatments that specifically target negative symptoms.

We and others have characterized subgroups of people with SRD that have a proportionally higher burden of negative symptoms, accompanied by markers of systemic inflammation (3–7). Recently, our group has focused on a sub-group defined by seropositivity for IgG anti-Gliadin Antibodies (AGA+), reflecting an immune reaction towards the gluten-derived protein gliadin that is found in wheat, barley, and rye (8, 9). This negative symptom predominant subgroup consists of about 32-38% of those with SRD. By contrast, fewer than 10% of non-diseased individuals are AGA+. The AGA+ SRD subgroup has repeatedly demonstrated elevated peripheral inflammatory proteins that are linked to both brain inflammation and negative symptom severity. When compared to AGA negative (AGA-) SRD, AGA+ SRD has robust elevations in the broadly pro-inflammatory cytokines including IL-1, TNF-α, and IL-6, as well as cytokines such as IFN-γ, IL-2, IL-4, IL-17, and IL-23 that are implicated in various T cell dysfunctions (5, 10, 11). Notably, other groups have described associations of Th17 and inflammatory cytokines with negative symptoms in SRD independent of AGA levels, suggesting that the association may generalize to other factors contributing to chronic inflammation in SRD (12–17).

Flow cytometry has revealed complex relationships between T cell abundance and negative symptoms in SRD. Elevated proportions of total T cells were associated with a higher burden of experiential negative symptoms across all SRD patients. By contrast, analyses of T cell subsets revealed that elevated proportions of CD4+ T cells and Regulatory T cells (Tregs) were associated with reduced experiential negative symptoms, but only in AGA+ SRD and not AGA-SRD (18). This divergent relationship between total T cell burden and T cell subsets, particularly the anti-inflammatory Treg population, suggests the involvement of an undetected pro-inflammatory T cell population contributing to negative symptom severity. Multiple activated T cell subsets could potentially play pathogenic roles, including Th17 cells, CD8+ cytotoxic T cells, mucosal-associated invariant T (MAIT) cells, natural killer T (NKT) cells, or other unconventional innate-like T cell populations.

The associations above have led to proposals to target inflammation to treat negative symptoms in SRD (19). Indeed, in two separate randomized controlled clinical trials, we found that a gluten free diet intervention in AGA+ SRD produced a modest reduction in experiential negative symptoms that coincided with a decrease in IL-17 (5, 10). These studies demonstrate that reducing sources of chronic inflammation can provide meaningful therapeutic benefit for SRD patients. However, targeting gluten may not benefit patients whose chronic inflammation arises from other factors, and gluten-free diets may be difficult to maintain in an outpatient setting. Detailed characterization of the underlying peripheral immune mechanisms would provide insight into specific immune states contributing to negative symptom psychopathology in SRD and aid in the identification of novel therapeutic targets.

Here, to address these questions, we performed single cell RNA-sequencing (scRNA-seq) on peripheral blood mononuclear cells (PBMCs) from 15 participants (SRD and healthy controls), enabling identification of immune cell subsets and characterization of their lineage, functionality, polarization state, migratory potential, and activation state (20, 21). In addition to detailed characterization of T cell populations, scRNA-seq allowed for parallel investigation of other immune cells (20, 21) including natural killer (NK) cells, monocytes, B cells, and dendritic cells (DC). We hypothesized that individuals with the AGA+ SRD would exhibit a distinct T cell-predominant inflammatory immunophenotype compared to AGA-SRD or Healthy Controls (HC), accompanied by enhanced inflammatory signatures among other immune cells such as NK cells, monocytes, B cells and/or DCs.

## Results

### A single-cell RNA sequencing atlas of peripheral immune cells in SRD subgroups

We performed single-cell RNA sequencing (scRNA-seq; Parse Biosciences Evercode WT) to characterize the transcriptional states of peripheral immune cells in SRD subgroups (Fig. 1A). Peripheral blood mononuclear cells (PBMCs) were collected from a subset of participants in a larger observational study of gluten sensitivity in SRD, including participants with SRD and high levels of AGA-IgG antibodies (≥20U, AGA+ SRD, N=7; Fig. 1B, Supplementary Table 1), participants with SRD and low AGA-IgG (N=3), and healthy controls with no history of mental illness (N=5). All five HC had AGA-IgG levels <20U. Within the SRD group, eight participants had a diagnosis of schizophrenia, one had a diagnosis of schizoaffective disorder bipolar type, and one had a diagnosis of schizoaffective disorder depressed type. Analyses of secondary psychiatric conditions and demographic variables revealed a low rate of comorbid conditions and no statistically significant differences in age, race, or sex between the three groups (Supplemental Table 2).

**Figure 1:**
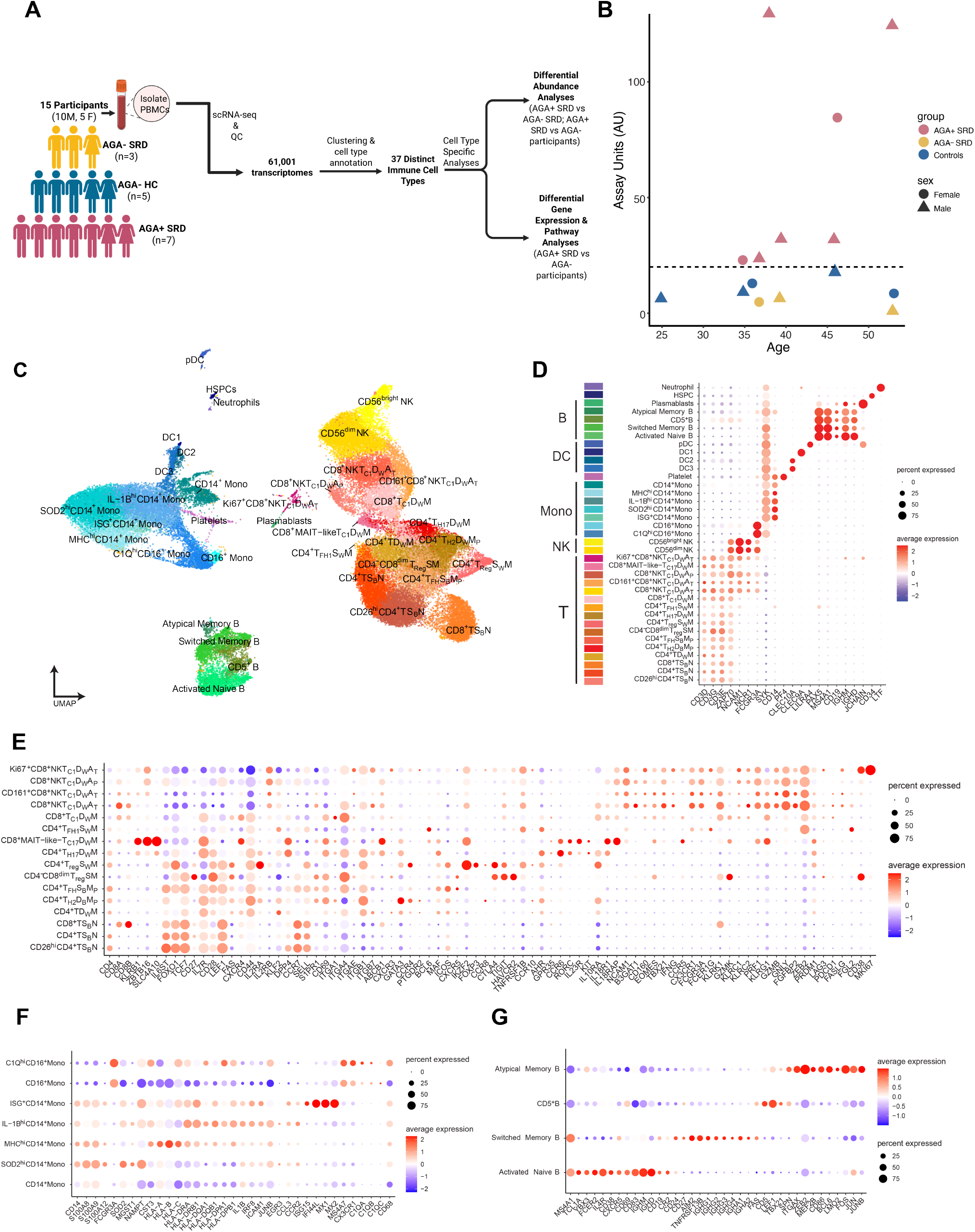
Study overview and identified immune cell types from scRNA-seq of PBMCs. **(A)** Overview schematic of study design. **(B)** Individual AGA-IgG level, age, and sex for each participant; Dashed line indicates AGA+ threshold of 20U. **(C)** UMAP of PBMCS clustered into 37 subpopulations: CD4^+^ T including naïve (CD4^+^TS_B_N, CD26^+^CD4^+^TS_B_N), memory (CD4^+^ TD_W_M), effector memory (CD4^+^T_H2_D_B_M_P_, CD4^+^T_FH_ S_B_M_P_, CD4^+^T_FH1_S_B_M, CD4^+^T_H17_D_W_M, CD4^+^T_reg_S_W_M), and effector stem-memory/memory precursor (CD4^+^T_H2_D_B_M_P_, CD4^+^T_FH_ S_B_M_P_); CD8+ T cells including naïve (CD8^+^ TS_B_N), memory cytotoxic-1 (CD8^+^ T_C1_ D_W_ M), terminally activated Natural Killer-T (NKT)-like effector (CD8^+^NKT_C1_ D_W_ A_t_, CD161^+^CD8^+^NKT_C1_D_W_ A_t_), memory precursor NKT-like effector cytotoxic-1 (CD8^+^NKT_C1_ D_W_ A_P_), proliferating terminally activated NKT-like effector cytotoxic-1 (Ki67^+^ CD8^+^NKT_C1_D_W_A_t_), Mucosal Associated Invariant T (MAIT)-Like cytotoxic-17 effector memory (CD8^+^MAIT-like-T_C17_D_W_M), and CD8-dimly positive regulatory memory precursor (CD4^-^CD8^dim^ T_Reg_ S_W_M_P_); CD14^+^ monocytes including classical (CD14^+^ Mono), (Major-Histocompatibility (MHC) gene-high (MHC^hi^ CD14^+^ Mono), superoxide dismutase 2 (SOD2)-high (SOD2^hi^ CD14^+^ Mono), IL-1B -high (IL-1B^hi^ CD14^+^ Mono), and interferon stimulated gene (ISG+ CD14^+^ Mono); CD16^+^ monocytes including non-classical (CD16^+^ mono) and Complement protein high (C1Q-High CD16^+^ Mono); B cells including activated Naïve, switched memory, atypical memory, CD5 positive (CD5^+^ B), and plasmablast (pb); dendritic cells (DC) including type 1 (DC1), type 2 (DC2), type 3 (DC3), and plasmacytoid (pDC); Natural Killer (NK) including CD56 brightly positive (NK CD56^bright^) and CD56 dimly positive (NK CD56^dim^); neutrophils; platelets; Hematopoietic Stem; Progenitor Cell (HSPCs). **(D)** Dot plot showing canonical marker genes used in the first level of annotation to identify broad immune cell lineages, with color bar indicating relative z-scored expression of each marker gene. **(E)** Dot plot showing marker genes used in the second level of annotation to T cell subpopulations, with color bar indicating relative z-scored expression of each marker gene. **(F)** Dot plot showing marker genes used in the second level of annotation for monocyte subpopulations, with color bar indicating relative z-scored expression of each marker gene. **(G)** Dot plot showing marker genes used in the second level of annotation for B cell subpopulations (excluding plasmablasts), with color bar indicating relative z-scored expression of each marker gene. Fig. 1A: Created in BioRender. Salem, D. (2026) https://BioRender.com/vha56u3

Unsupervised clustering of 61,001 high-quality cellular transcriptomes revealed 37 distinct immune cell clusters (Fig. 1C; Supplementary Fig 1). We annotated these to 16 T cell populations, two NK cell populations (CD56^dim^ NK and CD56^bright^ NK), seven monocyte populations, four dendritic cell populations (DC1, DC2, a monocyte like DC3, and plasmacytoid DC (pDC)), four B cell populations, plasmablasts, platelets, hematopoietic stem and progenitor cells (HSPCs) (Fig. 1D; Supplementary Fig 2-4; Supplemental Tables 3-7; Supplementary Note). Importantly, the expression patterns of canonical markers enabled us to assign each cluster of T cells, monocytes, and B cells to established functional subtypes (Fig. 1E-G).

**Figure 2:**
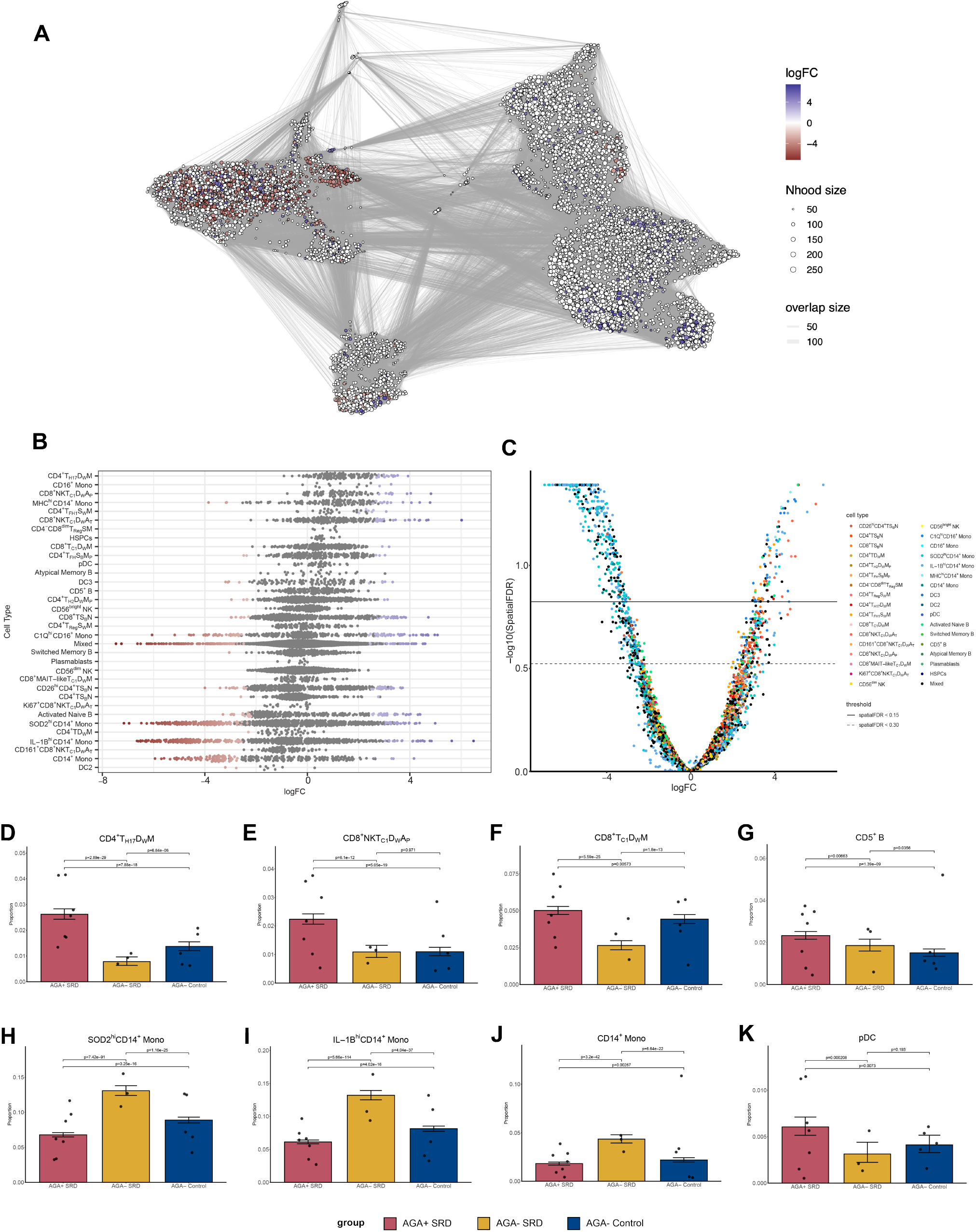
Differential abundance analyses show a higher predominance of T, B, and pDC immune subpopulations in AGA+ SRD and a lower abundance of monocyte subsets in AGA+ SRD. **(A)** Graph-based representation of neighborhoods identified by milo. Each node is a circle and depicts a neighborhood that is colored by their log fold change (if significant at FDR 30%***) comparing AGA+ SRD vs. AGA-participants (AGA-SRD plus AGA-HC). The size of each node depicts the neighborhood size (how many cells fall into that neighborhood) and edges depict the number of cells shared between adjacent neighborhoods. Non-differential abundance neighborhoods are colored white (FDR 30%). The layout of nodes corresponds to the position of the neighborhood index single cell in its respective UMAP embedding. **(B)** Beeswarm plot showing the distribution of log fold change in abundance between conditions in neighborhoods from different cell type clusters, with the most positive log fold change at the top of the plot and the most negative log fold change at the bottom of the plot. Positive log fold change indicates higher abundance in AGA+ SRD compared to AGA-participants, negative log fold change indicates lower abundance in AGA+ SRD compared to AGA-participants. Differential abundance neighborhoods are colored at FDR 30%. **(C)** Volcano plot showing each neighborhood’s respective log fold change and -log10(FDR) between AGA+ SRD compared to AGA-participants, with the color of each dot corresponding to a different immune cell subset. Solid line indicates FDR 15% and dashed line indicates FDR 30%. **(D) through (K)** Bar plots depicting pairwise comparisons using a binomial generalized linear model with logit link controlling for age and sex of the cell type proportion (%) of each immune cell subset in AGA+ SRD (red), AGA-SRD (yellow), and HC (blue), including **(D)** CD4^+^T_H17_D_W_M, **(E)** CD8^+^NKT_C1_D_W_ A_P_, **(F)** CD8^+^ T_C1_ D_W_ M, **(G)** CD5^+^ B, **(H)** SOD2^hi^ CD14^+^ Mono, (**I)** IL-1B^hi^ CD14^+^ Mono, **(J)** CD14^+^ Mono, and **(K)** pDC.

### AGA+ SRD is associated with increased abundance of T cell, B cell, and pDC sub-populations

We hypothesized that AGA+ SRD is associated with increased abundance of activated immune populations, compared to either AGA-SRD or healthy controls. Indeed, differential abundance analysis with Milo (22) revealed increased abundance (FDR < 0.3) of Th17 cells, Tfh1 cells, plasmacytoid dendritic cells (pDC), activated CD5+ B cells, and multiple CD8+ T-lineage populations (including NKT-like CD8+ cells, CD8+ effector/memory and naïve CD8+ T cells) (Figs. 2A-C). The strongest of these effects was for Th17 cells. Of the 120 identified Th17 cell neighborhoods, 94% were more abundant in AGA+ SRD, with a median log fold change of 1.37, including 16 neighborhoods significantly increased at FDR < 0.3. Pairwise contrasts using either Milo or a binomial generalized linear model confirmed significant enrichment of Th17 in AGA+ SRD relative to both comparison groups and a relative depletion in AGA-SRD (Fig. 2D; Supplementary Tables). Similarly, Tfh1, activated NKT-like and memory CD8+ Tc1 cells, CD5+ B cell, and pDCs were enriched in AGA+ SRD compared to both comparison groups (Figs. 2E-G). These results suggest a coordinated enrichment of pathogenic Th17 cells together with activated antigen-presenting and CD8+ T-lineage immune populations, consistent with a heightened and dysfunctional immune state in AGA+ SRD.

Parallel analyses also revealed a distinct immune signature in AGA-SRD, compared to AGA+ SRD and HC (FDR < 0.3; Figs. 2B-C). AGA-SRD was associated with increased abundance of several monocyte populations, including SOD2-Hi, IL-1B-Hi, and classical CD14^+^ monocytes, all of which exhibited a substantially increased abundance (25-35%) in AGA-SRD, compared to both comparison groups (Figs. 2H-J). These results suggest that AGA-SRD may be associated with immune signatures different from either AGA+ SRD or HC, although the very small sample size for AGA- SRD limited confidence in enrichments of specific cell populations.

We evaluated the specificity of immune signatures in AGA+ SRD vs. other conditions with elevated anti-gliadin antibodies by analyzing publicly available scRNA-seq of PBMC-derived T-cells from Celiac disease cases and controls (23). In contrast to AGA+ SRD, Celiac disease was not associated with substantial changes in the abundance of Th17 cells. The majority of Th17 cell neighborhoods were nominally depleted (log fold change < 0; Supplemental Fig. 5). These results provide evidence that the immune dysregulation in AGA+ SRD differs from that observed in Celiac disease.

### Dysregulation of metabolic and inflammatory signaling networks in AGA+ SRD

Next, we analyzed specific genes and pathways dysregulated in each cell type, comparing AGA+ SRD vs. AGA-SRD and HC combined via a conservative pseudobulk approach. Although few genes passed stringent multiple-testing thresholds, we detected 1,370 up-regulated and 2,677 down-regulated differentially expressed genes (DEGs) at a nominal p-value < 0.01, spanning 25 cell types (Fig. 3A; Supplementary Fig. 6).

**Figure 3:**
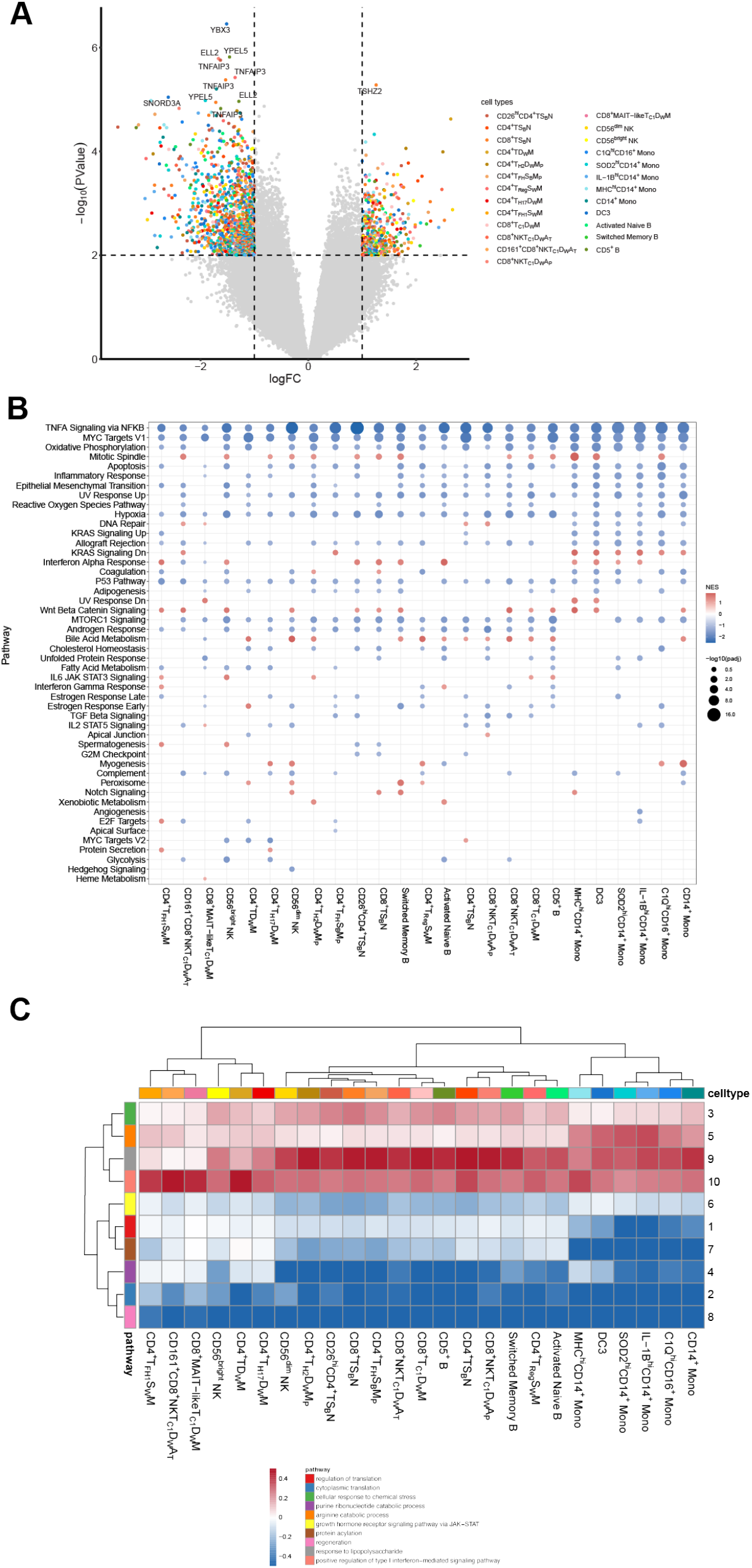
Differential gene expression and Gene Set Enrichment Analyses (GSEA) identify widespread dysfunction in several pathways in AGA+ SRD. **(A)** Volcano plot depicting the differential gene expression log fold change between AGA+ SRD and AGA-participants (AGA-SRD plus AGA-HC). Differentially expressed genes (DEGs) that are colored all have a log fold change magnitude of at least 1 and were nominally significant (p-value < 0.01). DEGs that passed multiple testing corrections are labeled. **(B)** Dot plot of normalized enrichment score (NES) from gene set enrichment analysis (GSEA) using DEGs in AGA+ SRD compared to AGA-participants. The size of each dot corresponds to the -log10(FDR-adjusted p-value) of each Molecular Signature Database (MSigDB) Hallmark pathway for select immune cell types. The color of each dot corresponds to the NES of the pathway for that immune cell type, with yellow indicating positive enrichment in AGA+ SRD, and dark blue indicating negative enrichment in AGA+ SRD. (**C**) Heatmap of K-means clustering of DEGs into immune transcriptional programs across immune cell types. The rows are annotated with the most overrepresented Gene Ontology biological process pathway for each module. The columns are annotated with immune cell types. Warmer colors represent modules upregulated in that cell type in AGA+ SRD compared to AGA-participants.

Gene set enrichment analyses (GSEA) of differential gene expression patterns in each cell type revealed dysregulation of multiple metabolic pathways in AGA+ SRD, spanning many immune cell populations (FDR < 0.05; Fig. 3B). These included pathways related to hypoxia, oxidative phosphorylation, glycolysis, and fatty acid metabolism, while bile acid metabolism was upregulated. GSEA also revealed a robust downregulation of TNF-α signaling via NF-kB, as well as of MYC target genes in all cell types.

By contrast, up-regulation of pro-inflammatory pathways occurred more selectively in specific immune cell types. These included protein secretion (Tfh1 and Th17), Interferon-alpha (IFN-α) response (Tfh1, CD8+ Tc1 memory, SOD2-Hi CD14+ mono, IL-1B-Hi CD14+ mono, and DC3), WNT/β-catenin signaling (CD4+ T memory, Tfh1, CD8+ NKTc1, CD5+, and classical CD14+ mono), and IL-6 JAK/STAT3 signaling (Tfh1 and CD5+ B) (Fig. 3B). These findings suggest a shift from an acute and innate inflammatory process such as one driven by TNF-α, to a chronic inflammatory process with metabolic changes that may facilitate cell survival, subtype maturation, and cytokine responsiveness.

K-means clustering of DEG log fold changes (k=10) provided further insights into these coordinated transcriptional changes across cell types (Fig. 3C; Supplementary Fig. 6). Two modules were upregulated in distinct populations of activated T-cells from AGA+ SRD. Module 6 was up-regulated in Tfh1, Th17, MAIT-like, CD161+ NKT, and memory CD4+ T-cell populations and was enriched for genes involved in growth hormone receptor signaling through the JAK/STAT pathway. Module 10 was upregulated in Th2, Th17, Tfh, CD8+ NKT, and CD8+ Tc1 populations but displayed minimal changes in Tfh1, CD161+ CD8+ NKT Tc1, and CD8+ MAIT-like cells; this module was enriched for genes involved in positive regulation of type I interferon signaling. These results suggest that multiple cytokines contribute to the activation states in AGA+ SRD, with varying responsiveness across T-cell subtypes.

Other modules displayed transcriptional dynamics primarily within myeloid populations. Module 5 was down-regulated in several myeloid populations and was enriched for genes involved in the arginine catabolic process, suggesting dysregulation of immunometabolic pathways. Module 9 was selectively downregulated in monocyte populations and was enriched for genes involved in the response to lipopolysaccharide (LPS), a central mediator of innate immune activation, consistent with dysregulated inflammatory signaling in monocytes from AGA+ SRD.

Together, these findings support widespread remodeling of innate immune and inflammatory signaling pathways in AGA+ SRD, including prominent cell type-specific alterations in JAK/STAT and interferon signaling, as well as monocyte inflammatory states.

## Discussion

To our knowledge, this is the first study investigating the transcriptomes of peripheral immune cells at a single-cell resolution in SRD. Our results suggest that AGA+ SRD is associated with an increased abundance of effector/activated T and B cell subpopulations (adaptive immune cell types), while AGA-SRD was associated with increased abundance of monocyte subpopulations (innate immune cell types). Differential gene expression and network analyses identified multiple signaling pathways that may drive changes within specific cell types, including JAK/STAT, WNT/β-catenin, and IL-6 signaling in distinct T cell groups.

In line with our prior finding of an elevated Th17 cell cytokine signature (IL-6, IL-17, GM-CSF) in AGA+ SRD (24), our most significant cell type enrichment in AGA+ SRD was in Th17 cells, which are well-established drivers of autoimmunity that preferentially persist and promote chronic inflammation by secreting IL-17 in an IL-6-and IL-23-dependent manner (25). In multiple sclerosis, IL-17 secreted from Th17 cells was found to weaken the blood-brain barrier (BBB) by binding to IL-17R on endothelial cells, leading to easier central nervous system (CNS) infiltration by Th17 cells via CCR6 and VLA-4 (26, 27). Our findings of increased expression of pathogenic markers, including *IL23R, CCR6, KLRB1*, and *ABCB1*, and high expression of CNS migration markers in *ITGA4* and *ITGB7* (the genes coding for the integrin VLA-4) suggest CNS pathogenicity potential in AGA+ SRD (27–29). Functionally, Th17 cells in AGA+ SRD also have upregulated JAK/STAT (a driver IL-17/IL-23 inflammatory axis) (30) and protein secretion pathways, also supporting pathogenicity potential.

In addition to Th17 cells, we identified two cell types with enriched IL-6/STAT3 signaling that were more abundant in AGA+ SRD: CD5+ B cells (which are a known contributor in systemic IL-6 production) (31) and Tfh1 cells. CD5 is upregulated on activated B cells that have the capacity to differentiate into auto-antibody-secreting plasma cells, as seen in lupus, where CD5+ B cell populations are similarly abundant (32). The CD5+ B cell cluster was enriched in *TBX21* expression (coding for T-bet), a marker associated with atypical, germinal center maturation, and IgG2 skewed antibody secretion that can result in enhanced NK and NK-like CD8+ T cell-mediated antibody-dependent cell toxicity (33). It is possible that the Tfh1 cells are promoting CD5+ B cell maturation and effector capacity, as they can directly expand T-bet+ B cells via IFN-γ secretion (33, 34).

We also found a higher abundance of several CD8+ T cell subsets in AGA+ SRD, most pertinently in effector/memory, effector progenitor, and terminal effector cell populations. These CD8+ T cells exhibited a Tc1 polarization, and the effector populations expressed NKT markers, indicating high activation, cytotoxic potential, and IFN-γ secretion, which is increased in the AGA+ SRD subgroup compared to AGA- SRD. Th17 cells are known to augment CD8+ T cell activity by enhancing their migratory potential, expanding cytotoxic NK-like T cell populations, and by interfering with Treg suppressive function (35, 36). We have previously found Tregs to be protective of negative symptoms, while another, unspecified T cell population (potentially CD8+ T cells) was found to be associated with worse negative symptom severity in AGA+ SRD (18). It is possible that an imbalance of Th17 cells and Tregs could be driving this expansion of cytotoxic effector CD8+ T cells that may be implicated in the negative symptoms seen in AGA+ SRD. Our pathway analyses are suggestive of this, with upregulated bile acid metabolism (a pathway known to affect Treg/Th17 balance) and WNT/β-catenin (a driver of Th17 inflammation) pathways in several T cell subpopulations and upregulated JAK/STAT and IL-6 pathways that can drive Th17 inflammation (37, 38).

Interestingly, we also found a higher abundance of pDCs in AGA+ SRD. pDCs normally mount robust anti-viral responses, are amongst the strongest producers of type I interferons (type-1 IFN, including IFN-α and IFN-β), and are potent CD8+ T cell activators (39). pDCs have been implicated in autoimmunity after responding to DNA, such as from cells destroyed by CD8+ T cell inflammation, and activation of pDCs could further prime CD8+ T cells to target self-antigens and further perpetuate their dysfunction (39). Of note, pDCs have not been extensively studied in SRD to date, largely due to their relatively lower cell counts, but IFN-α has been found to be elevated in SRD, suggestive of pDC activity (40). Type I IFN therapies for viral infections characteristically produce deficits in anhedonia and motivation that are clinically similar to experiential negative symptoms, so our finding that pDCs are increased in abundance may be related to the correlation between negative symptoms and inflammation we see in AGA+ SRD (41).

Notably, there is a decreased abundance in several monocyte populations in AGA+ SRD compared to AGA-participants. These monocyte subsets, including SOD2-Hi CD14+ monocytes, IL-1B-Hi CD14+ monocytes, and classical CD14+ monocytes, all generally had significant downregulation of pathways related to cell proliferation, such as cytoplasmic translation, regulation of translation, and KRAS signaling. The decreased abundance of monocytes in AGA+ SRD may be due to cellular dysfunction from chronic inflammation and decreased cell production, rather than increased cell death, as apoptosis and p53 pathways were downregulated in AGA+ SRD. It is also possible that our AGA- SRD population is enriched in a monocyte predominant inflammatory phenotype, as we found in pairwise differential abundance comparisons that AGA-SRD had a higher abundance of several monocyte subsets compared to either AGA+ SRD or HC. Due to the small sample size, we were underpowered to detect differences in AGA-SRD compared to either HC or AGA+ SRD in DEG and pathway analyses, which limited our ability to ascertain functional differences in monocytes in AGA-SRD.

We note several limitations of the current study. The small sample size impacted our ability to detect DEGs with more moderate effect sizes. A larger sample size would also enable stratified analyses by sex, antipsychotic medication, or presence of comorbid inflammatory conditions (e.g., HIV (42)). Experimental studies of *ex vivo* immune cells could directly test the functional states of populations dysregulated in AGA+ SRD, as well as causal roles for intra- and intercellular signaling pathways.

The lymphocyte predominant inflammatory signature in AGA+ SRD suggests an antigen-antibody mediated process specific to gliadin, potentially driven by CD5+ B cells and interactions between pro-inflammatory T cell subsets (i.e. Th17, CD8+ T cell, and Tfh1 cells) with downstream effects on BBB integrity, neuroinflammation, and negative symptom pathophysiology (Fig. 4) (43). This model is further supported by two independent randomized control clinical trials investigating the efficacy of a gluten free diet intervention in AGA+ SRD for negative symptoms, which resulted in a reduction in experiential negative symptoms and decreases in Th17-related cytokines, most notably IL-2, IL-17, and IL-23 (5, 10).

**Figure 4:**
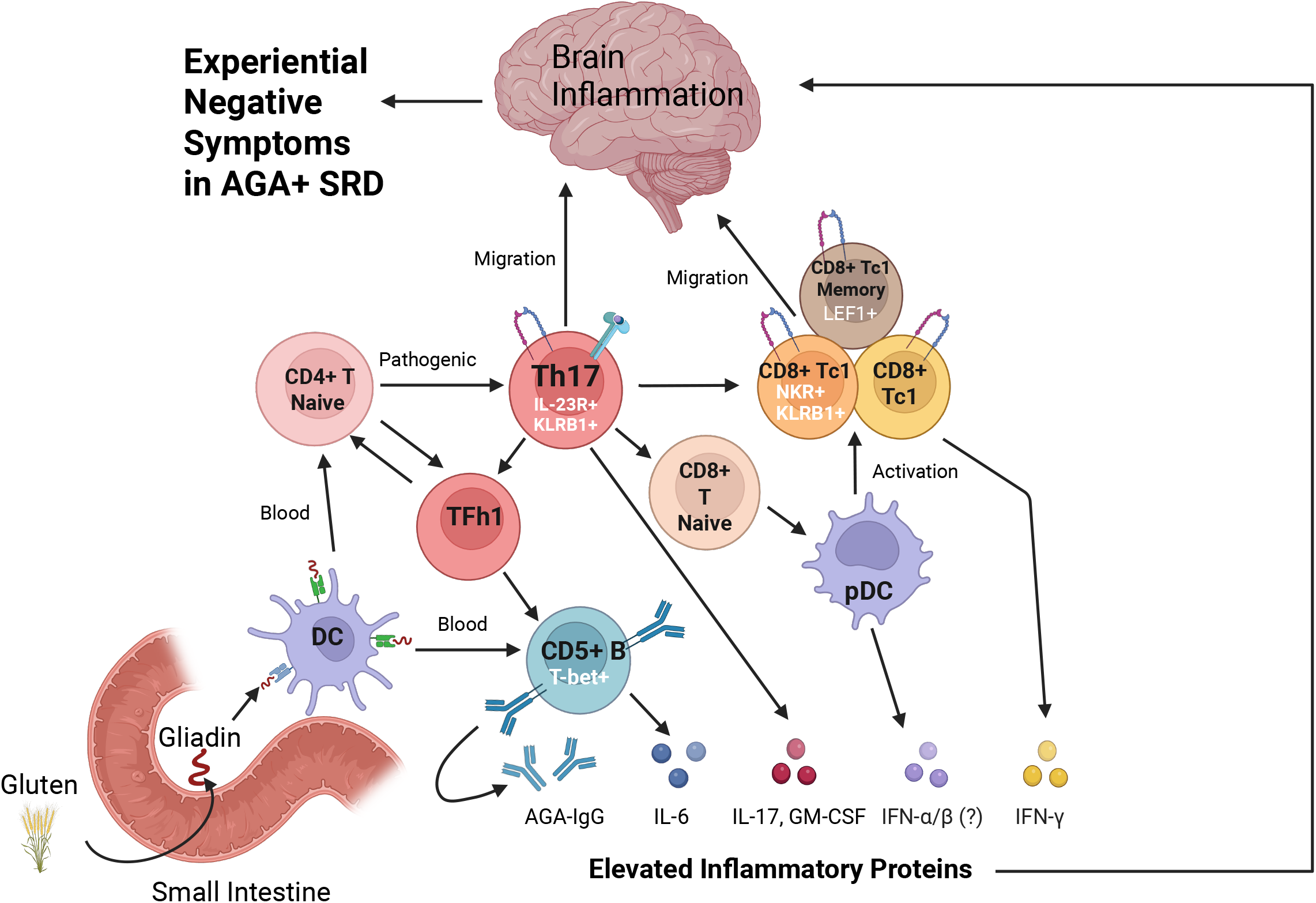
Conceptual overview of T cell predominant inflammation and potential mechanism in the pathogenesis of experiential negative symptoms in AGA+ SRD. Gluten ingestion and subsequent digestion in the small intestine gliadin elicits an inappropriate local inflammatory response that subsequently leads to systemic inflammation. Dendritic Cells (DCs), as primary antigen presenting cells, present gliadin to B cells, which become activated and express CD5, subsequently gaining the ability to secrete AGA-IgG. In tandem, DCs present gliadin antigen to naïve CD4+ T cells which differentiate into Tfh1 cells. Tfh1 cells can provide help for the maturation of effector CD5+ B cells with upregulation of T-bet and IL-6 secretory function. Additionally, TFH1 cells along with the higher IL-6 levels together can help differentiation of Th17 cells expressing pathogenic markers such as IL-23R and KLRB1. Th17 cells, through secretion of IL-17 and GM-CSF, can recruit and increase the cytotoxicity and effector potential of CD8+ T cells and can attract naïve CD8+ T cells, which with the help of plasmacytoid dendritic cells (pDCs), can also help with CD8+ T cell maturation into effector CD8+ T cells expressing Natural Killer Receptors, KLRB1, and stem-like markers including LEF1. Additionally, IL-17 secreted by TH17 cells can increase blood-brain permeability, allowing for easier brain migration of central nervous system (CNS) homing effector T cells that express ITGA4 and ITGB7, including the CD8+ T effector cells and pathogenic TH17 cells. IFN-G secreted by effector CD8+ TC1 cells, with GM-CSF and potentially Type 1 Interferons, can lead to more immune dysfunction contributing to CNS inflammation. With the increased brain inflammation in AGA+ SRD, central processes such as neurotransmitter changes and downstream functional connectivity changes downstream can contribute to the pathogenesis of experiential negative symptoms (anhedonia and motivational deficits) that are chronic in SRD. Created in BioRender. Salem, D. (2026) https://BioRender.com/6ssdrt4

Thus, our main finding of a T cell predominant inflammatory phenotype in AGA+ SRD, particularly one enriched in Th17 inflammation, has enormous clinical implications for future work. Currently, the standard of care in treating negative symptoms in SRD focuses on not exacerbating them (such as through high-potency D2 antagonist antipsychotics and prioritizing the use of clozapine), rather than on specifically decreasing their severity (44). In identifying the underlying biological dysfunction in this negative symptom predominant AGA+ SRD subgroup, future clinical trials can focus on targeting this specific immune dysfunction by using specific Th17 depleting agents such as IL-23 and IL-17 inhibitors. Similarly, as we found upregulated JAK-STAT pathway activity, the use of JAK-inhibitors may be another avenue to target in future work (45). Overall, our findings support the role of AGA-IgG positivity as a biomarker in identifying a distinct Th17 inflammatory subgroup within SRD. Future studies with larger sample sizes to substantiate the distinct immune signatures we identified in SRD are warranted.

## Methods

### Study participants

In this cross-sectional study consisting of a one-time peripheral blood draw, we initially recruited a total of 12 participants with SRD and 6 HC from the Maryland Psychiatric Research Center, University of Maryland School of Medicine. Participants were of either sex or race were included if they were between the ages of 18 and 64 years old (Supplemental Table 1). Participants with SRD were required to meet Diagnostic and Statistical Manual (DSM)-5 criteria for either schizophrenia or schizoaffective disorder. HC participants were included if they were free of all major psychiatric illnesses as per the DSM-5. As a part of the AGA-IgG analysis, before assaying AGA-IgG levels, all participants must be negative for the celiac disease marker anti-tissue transglutaminase and thus were excluded if positive (presumed celiac disease). As a part of scRNA-seq quality control, participants with overall low-quality cells were excluded from further analysis. The study was approved by the University of Maryland Institutional Review Board (IRB). All participants were required to provide written informed consent and sign the Evaluation to Sign Consent to assess capacity.

### Sample and demographics collection and AGA-IgG level assessment

All study participants arrived between 0700 and 0800 under fasting conditions. Participants had their blood drawn for assessment of AGA-IgG levels and for scRNA-seq analysis. We measured AGA-IgG levels at the Institute for Genome Sciences using the Werfen Diagnostics kit catalog #708655. Units were determined by the manufacturer based on a semiquantitative detection of Gliadin IgG antibodies in human serum. AGA+ status was defined as ≥20U AGA-IgG and AGA-status was defined as <20U AGA-IgG based on manufacturer normative ranges and our prior work (4, 5, 9– 11). Additionally, demographic information, including age, sex, and race, was collected. For sex and race, we utilized descriptive statistics and compared differences between AGA+ SRD, AGA-SRD, and HC using Fisher’s Exact Test. For age and AGA-IgG, we utilized a one-way ANOVA to assess group differences between AGA+ SRD, AGA-SRD, and HC (Supplemental Table 2).

### Single Cell RNA-sequencing and Quality Control

Single cell RNA sequencing (sc-RNA-seq) cell annotation was performed using the Parse Biosciences Evercode WT platform on 18 participants. After integration, there were 74,273 cells in total. Cells were filtered out if the percentage of mitochondrial DNA was greater than 15% and/or if the number of unique molecular identifiers (UMIs) was lower than 500 or greater than 10,000. Three participants (2 SRD, 1 HC) were removed due to overall low-quality cells. Cells from the remaining 15 participants (n=10 SRD, n=5 HC) were next subset by participant, and then doublets were detected using the package DoubletFinder (46), which simulated artificial doublets by combining the expression profiles of single-cell data in different clusters. After quality control, 61,001 transcriptomes remained from 15 total participants. Integration local inverse Simpson’s Index (iLISI) and cell type local inverse Simpson’s Index (cLISI) scores were calculated using the lisi package (47) to measure the quality of integration and clustering (Supplemental Fig. 1).

### Cell Type Annotations

Following quality control and post-integration, we performed unsupervised clustering of cells into groups of cells with similar expression profiles using Seurat V5 (48), which revealed 37 distinct immune cell clusters. Clusters were annotated first at a broader immune cell level using canonical marker genes to identify 16 T cell clusters, 2 NK cell clusters, 7 monocyte clusters, 1 platelet cluster, 4 DC cluster, 5 B lineage (including 4 B cell clusters and 1 plasmablast), and 1 Hematopoietic Stem and Progenitor Cell (HSPCs) cluster (Figure 1D). T cell clusters were identified as strongly expressing *CD3D, CD3G, CD3E*, and *ZAP70* (21, 49). NK cell clusters were identified as strongly expressing *NCAM1* and *NCR1* while having low expression of *CD3D, CD3G, or CD3E* (50, 51). Monocytes were identified by having strong expression of *SYK* and *CD14* and/or *FCGR3A (52, 53)*. Platelets were identified as having strong expression of *PF4* (54). DC clusters were identified by strong expression of *SYK* and *CLEC10A, CLEC9A*, or *LILRA4* (55–57). B cell clusters were identified by strong expression of *SYK, PAX5, MS4A1, CD19, IGHM, IGHD* and low expression of *JCHAIN*. Plasmablasts were identified as having positivity for *IGHM, IGHD*, and having the strongest expression for *JCHAIN* (62). HSPCs were identified by strong expression of *CD34* (63). Neutrophils were identified by strong expression of *LTF* (64).

In the next layer of annotation, we further annotated each broad immune cell cluster that had more than one identified cluster, including the T cells, NK cells, monocytes, DCs, and B cells. The goal of this deeper secondary level of annotation for all clusters was to identify characteristic markers of lineage, function, migratory potential, and activation status. To achieve this, we performed a comprehensive literature search to identify known key markers of known immune cell subsets and utilized the Seurat FindAllMarkers function (48). DC clusters were further annotated into Dendritic Cell type 1 (DC1), Dendritic Cell type 2 (DC2), Dendritic Cell type 3 (DC3), and Plasmacytoid Dendritic Cell (pDC) (Supplemental Table 3). B cell clusters were annotated further by assessing the expression of markers related to maturation stage, antibody class isotype, proliferation, and atypical markers (Supplemental Table 4). NK cells were further annotated into the two broad lineages of CD56+dim NK cells and CD56+bright NK cells using the relative expression of *NCAM1* (Supplemental Table 5). Monocytes were further annotated into broad lineage based on expression of *CD14* and/or *CD16*, followed by other major subtypes identified from these monocyte immune atlases (50, 63–67), assessing for expression of markers related to antigen expression, oxidative stress, IL-1B expression, Interferon Stimulation Gene (ISG) expression, and *C1Q* expression (Supplemental Table 6).

To annotate T cell clusters, we utilized the modular nomenclature framework detailed in the 2026 T cell nomenclature consensus statement, which allowed us to concisely annotate T cell clusters based on lineage (CD4+ T, CD8+ T, NKT, and MAIT-like), function (T-helper (Th)1, Th2, Th17, Treg, T-cytotoxic (Tc) 1, Tc2, Tc17), migration potential, and differentiation state (21) (Supplemental Table 7). Migration potential for T cells was annotated first using the two major headings based on the expression of *CD62L* and *CCR7*: “S” for secondary lymphoid organ (SLO)/uninflamed lymph node migration potential indicated with strong expression of *CD62L* and *CCR7*, or “D”, for disseminated status (cells that do not tend to migrate to uninflamed SLOs/lymph nodes) indicated by low expression of *CD62L* and *CCR7*. These migratory major headings of S and D were then further annotated with subscripts of “W”, indicating widespread recirculatory migration potential between blood and nonlymphoid tissues (for example, a T cell annotated with “D_W_” indicates the potential to migrate to the gut and circulate in the blood) or “B” for being found in blood (for example, a T cell annotated with “S_B_” indicates SLO migratory potential but is found in the blood). Differentiation state for T cells was annotated using the following major headings: “N” for naïve, “A” for activated, and “M” for memory. Differentiation state subscripts were further added when applicable, including the subscript of “p” for progenitor/precursor or “t” for terminally differentiated.

### Differential Abundance Testing

The KNN graph was built using a K-value of 45 and a d-value of 20. Multiple contrasts were used to capture the differences in cell type abundance, including AGA+ SRD vs. AGA-participants, AGA+ SRD vs. AGA-SRD, AGA+ SRD vs. AGA-controls, AGA-SRD vs. AGA-controls, and SRD vs. HC. The model included the group, sex, and age, as well as the contrast of interest for statistical testing. The neighborhoods were annotated using the cell type labels that were manually curated during the annotation step, and any neighborhoods that did not have at least a 70% majority of a single cell type were then labeled as “Mixed.” The neighborhoods were grouped by their cell type annotation, and then the median log fold change was calculated for each cell type.

### Differential Gene Expression Analysis

The scRNA-seq counts were pseudobulked by sample and cell type, and differential gene expression was tested using the package edgeR (70). Gene-level counts were modeled using negative binomial generalized linear models (GLMs), with normalization factors calculated using the trimmed mean of M-values (TMM) method. Dispersion parameters were estimated using empirical Bayes methods, allowing for gene-specific variability while borrowing information across genes to improve stability. A quasi-likelihood framework was used to fit the GLMs, and significance testing was done with quasi-likelihood F-tests. The Benjamini-Hochberg false discovery rate (FDR) was used to adjust the p-values for multiple testing.

### Gene Set Enrichment Analysis

The differential gene expression results were used to create a ranked list for each cell type, which was then used as input for gene set enrichment analysis using the package fgsea (71). Hallmark gene sets with a minimum of 15 genes and a maximum of 500 genes from the Molecular Signatures Database (MSigDB) were used as the pathways tested. Significance testing was permutation-based (*N* = 10000), and the results were adjusted using the Benjamini-Hochberg correction.

### K-Means Clustering of Gene Modules

The differential gene expression results were also used for k-means clustering to identify modules with similar patterns of expression between cell types. A K of ten was used, and each module was tested using Gene Ontology biological processes to identify the most overrepresented term within each gene module.

## Notes

### Competing Interest Statement

The authors have declared no competing interest.

