## Supplemental Material for "Molecular adaptations of activated T-cells in an inflammation-associated schizophrenia sub-group"

### Supplemental Materials

#### Supplementary Note

##### *Cell Annotation:*

We next performed a secondary level of annotation of the broad immune lineages that had multiple clusters, including the NK cells, monocytes, DCs, B cells, and T cells. The complete list of markers used in this secondary level of annotation can be found in Supplementary Tables 3-7.

Of the NK cell clusters, we identified one cluster that had higher expression of *NCAM1* which codes for CD56, which is canonically used to differentiate between a CD56 "dim" cytotoxic and pro-inflammatory phenotype of NK cells and a CD56 "bright" NK cell that is more regulatory in phenotype. Consistent with this, we found a higher expression of *FCGR3A, GZMB, CX3CR1, LILRB1,* and *B3GAT1* suggesting a mature and cytotoxic phenotype in the NK CD56-bright cluster compared to the NK CD56-dim cluster, which had higher expression of *GZMK, KLRC1, KLRC2, IL7R, and SELL*, indicating a comparatively regulatory and immature phenotype (Supplemental Fig. 4; Supplemental Table 5).

Among the four B cell clusters, we found one cluster of naïve B cells that had characteristically low expression of IgG or IgA related genes and had the strongest expression of several Naïve B cell markers, including *PAX5, IGHD, IGHM, IL4R, TCL1A, YBX3, and FCER2*. Notably, this naïve B cell cluster also had strong expression of the germinal center homing chemokine gene *CXCR5* and strong expression of the activation markers *CD69* and *CD83*, together suggesting an activating and expanding naïve B cell population. Two B cell clusters had strong expression of memory markers. The first memory B cell cluster characteristically had strong expression of *CD27* and had the strongest expression of IgG and IgA related genes, altogether indicating a class-switched memory B cell phenotype. The second memory B cell cluster was identified to be an "atypical" (also known as "double negative") B cell phenotype, with low expression of the typical memory marker *CD27*, and low expression of naïve B cell markers including *IGHD* and *CR2*. This atypical memory B cell cluster had strongest expression of *ITGAX* (which encodes the protein CD11c) and had positive expression of *TBX21*, which is a phenotype of B cells seen to be expanded in autoimmune conditions and in aging. The last B cell cluster had strong expression of *CD27, SPN,* and *CD5*. CD5 positivity is seen in activated B cell subsets, most notably plasmablast precursor B cells and B1-B cells. (Fig. 1G; Supplemental Table 4).

In the 4 DC clusters, we found one which had the strongest expression of *CLEC9A* and *XCR1*, while being negative for expression of *LILRA4*, indicating a dendritic cell type 1 (DC1), which are antigen presenting cells that preferentially activate CD8+ T cells. There was one DC cluster that had strong expression of *LILRA4*, *CLEC4C,* and lymphoid markers (*JCHAIN* and *TCF4)*, which was consistent with a lymphoid derived plasmacytoid dendritic cell (pDC) lineage. The remaining 2 DC clusters both had strong expression of *CLEC10A* and *ITGAX* but were distinguishable by one cluster having strong expression of *CD5* and low expression of CD14 consistent with a dendritic cell type 2 (DC2) phenotype, while the other cluster was negative for CD5 but strongly expressed *CD14* and *CD163*, consistent with a dendritic cell type 3 (DC3) phenotype (Supplemental Fig. 3; Supplemental Table 3). Notably, DC2 and DC3 both present antigen to helper T cells, but DC3 preferentially elicits a Th17 related response, while the CD5+ DC2 phenotype more broadly polarize to Th17, Treg, Th2, or Th22.

The 7 monocyte clusters were then assessed for the expression of *CD14* and/or *FCGR3A*, which revealed 5 clusters of CD14+CD16- monocytes and 2 clusters of CD14-CD16+ monocytes. Of the 5 CD14+ monocyte clusters, we identified one cluster which fit a classical monocyte signature (CD14+ mono), one cluster which had the highest expression of Major Histocompatibility Complex (MHC) I and II gene (MHC-Hi CD14+ mono) indicating robust antigen presentation capability, one cluster which had the highest expression redox related genes (SOD2-Hi CD14+ mono), one cluster with the highest expression of *IL1B* and migratory genes *CCR2* and *CCL3* (IL-1B-Hi CD14+ mono), and the last cluster which had high expression of interferon related genes *ISG15, IFI44L, MX1,* and *MX2* (ISG+ CD14+ mono). In the two CD16+ monocyte clusters, we found high expression of complement protein genes *C1QA* and *C1QB* and macrophage differentiation gene *CD68* to be brightest in one cluster (C1Q-Hi CD16+ mono) indicating a more developed and pro-inflammatory monocyte population compared to the other CD16+ mono cluster which appeared comparatively less differentiated (Fig. 1F; Supplemental Table 6).

We assessed the 16 T cell clusters for the expression of *CD4, CD8A*, and *CD8B*, finding 8 CD4+ T cell clusters and 8 CD8+ T cell clusters (Fig. 1E; Supplemental Table 7). Among the 8 CD4+ T cell clusters, we found 2 naïve CD4+ T cell clusters that both had strong expression of secondary lymphoid organ (SLO) “central” homing markers *CD62L* and *SELL*, with one of the clusters (CD26$^+$ CD4$^+$ TS$_B$N) having stronger expression of *DPP4* and *ICOS,* indicating a relatively higher differentiation state. The remaining 6 naïve CD4+ T cell clusters had markers indicative of a memory phenotype, with one cluster being non-polarized, one cluster having strong expression of *GATA2* and *PTGDR2* indicating a T-helper-2 (Th2) polarization, one cluster having strong expression of *RORC, CCR6,* and *IL23R* indicating a T-helper-17 (Th17) polarization, one cluster having strong expression of *FOXP3* and *IL2RA* indicating a Treg polarization, one cluster having strong expression of *CXCR5* indicating a Tfh (T-follicular-helper) polarization, and the final cluster having strong expression of *BCL6* and *TBX21* indicating a Tfh1 (T-follicular-helper-1) polarization. The Tfh and Th2 clusters additionally had stronger expression of naïve markers *FAS*, *LEF1*, *IL6ST* and *TCF7* in addition to their positivity for memory markers (*CD44, CD69)*, indicating the presence of memory progenitor phenotypes. We also assessed migration potential in each of these CD4+ T cell clusters, finding widespread tissue migratory potential in the non-polarized (CD4$^+$ TD$_W$M) and Th17 clusters (CD4$^+$ T$_{H17}$D$_W$M), non-SLO migratory potential but possible tissue homing potential in the Th2 cluster (CD4$^+$ T$_{H2}$D$_B$M$_p$), SLO migratory potential in the Tfh (CD4$^+$ T$_{FH}$S$_B$M) and Tfh1 (CD4$^+$ T$_{FH1}$S$_W$M) clusters, and widespread SLO and uninflamed tissue migratory potential in the Treg (CD4$^+$ T$_{reg}$S$_W$M) cluster. Of note, the Th17 cluster had expression of the innate-like markers *KLRB1, ZBTB16,* and *SLC4A10.*

Among the 8 CD8+ T cell clusters, we found 4 with strong expression of Natural Killer T (NKT) markers that had a cytotoxic-1 (Tc1) polarization, one memory cluster with strong expression of Mucosal Associated Invariant T (MAIT)-like markers with a cytotoxic-17 (Tc17) polarization and widespread uninflamed tissue migratory potential, one memory CD8+ T cell cluster with widespread uninflamed tissue migratory potential, one naïve CD8+ T cell cluster with a strong expression of SLO homing markers (CD8$^+$ TS$_B$N), and one memory precursor CD8-dim T cell cluster with a regulatory polarization and with widespread SLO and uninflamed tissue migratory potential. All 4 of the CD8+ NKT-like clusters had widespread uninflamed tissue migratory potential and had expression of characteristic natural killer markers (*NCAM1, CD160, FCGR3A)* with one cluster additionally having high expression of the proliferation marker *MKI67* and one other cluster having higher expression of *KLRB1* (CD161+), a marker more common in MAIT-like populations. Additionally, all 3 of the NKT-like CD8+ Tc1 clusters expressed activation markers strongly (*KLRG1, CX3CR1, TBX21, B3GAT1)* consistent with a terminally activated differentiation state (A$_t$) and one cluster additionally having high *LEF1* expression supporting a long-lived/effector progenitor (A$_p$) differentiation state.

### Supplementary Figures:

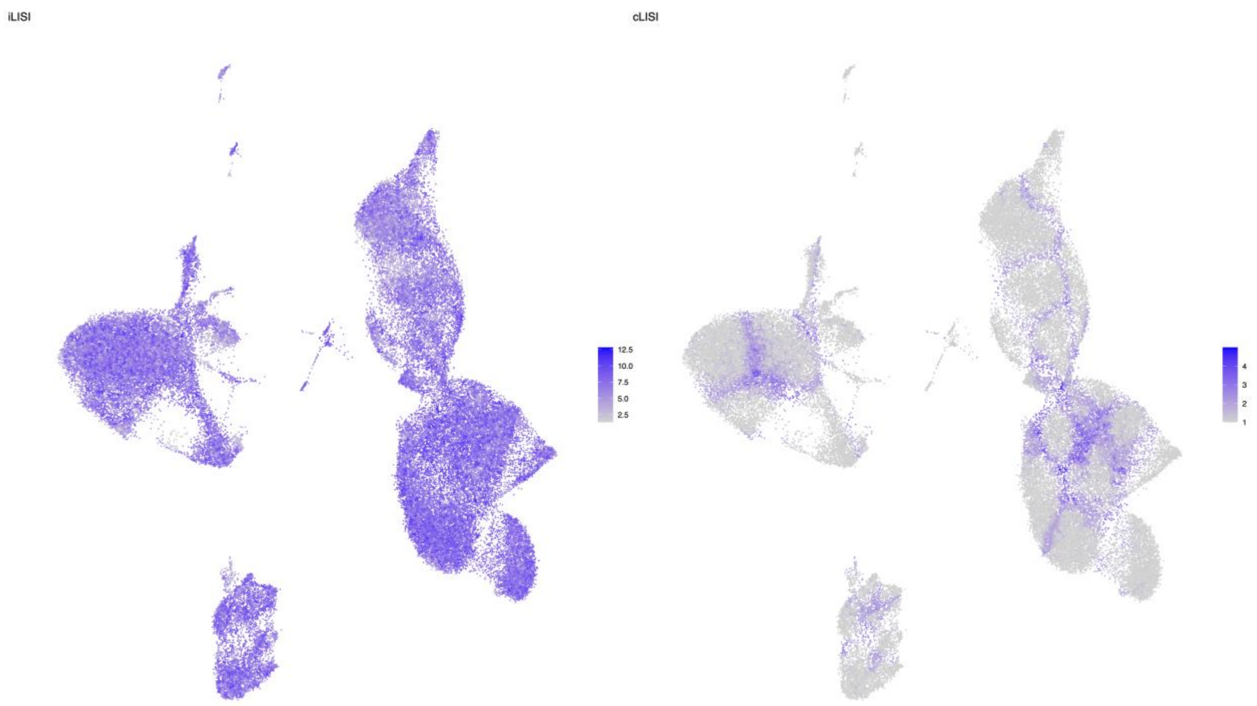


**Supplemental Figure 1: UMAP colored by LISI Scores.** Integration local inverse Simpson's Index (iLISI) and cell-type local inverse Simpson's Index (cLISI) scores are shown on the UMAP, representing how well integrated the data is and the quality of the clustering, respectively.

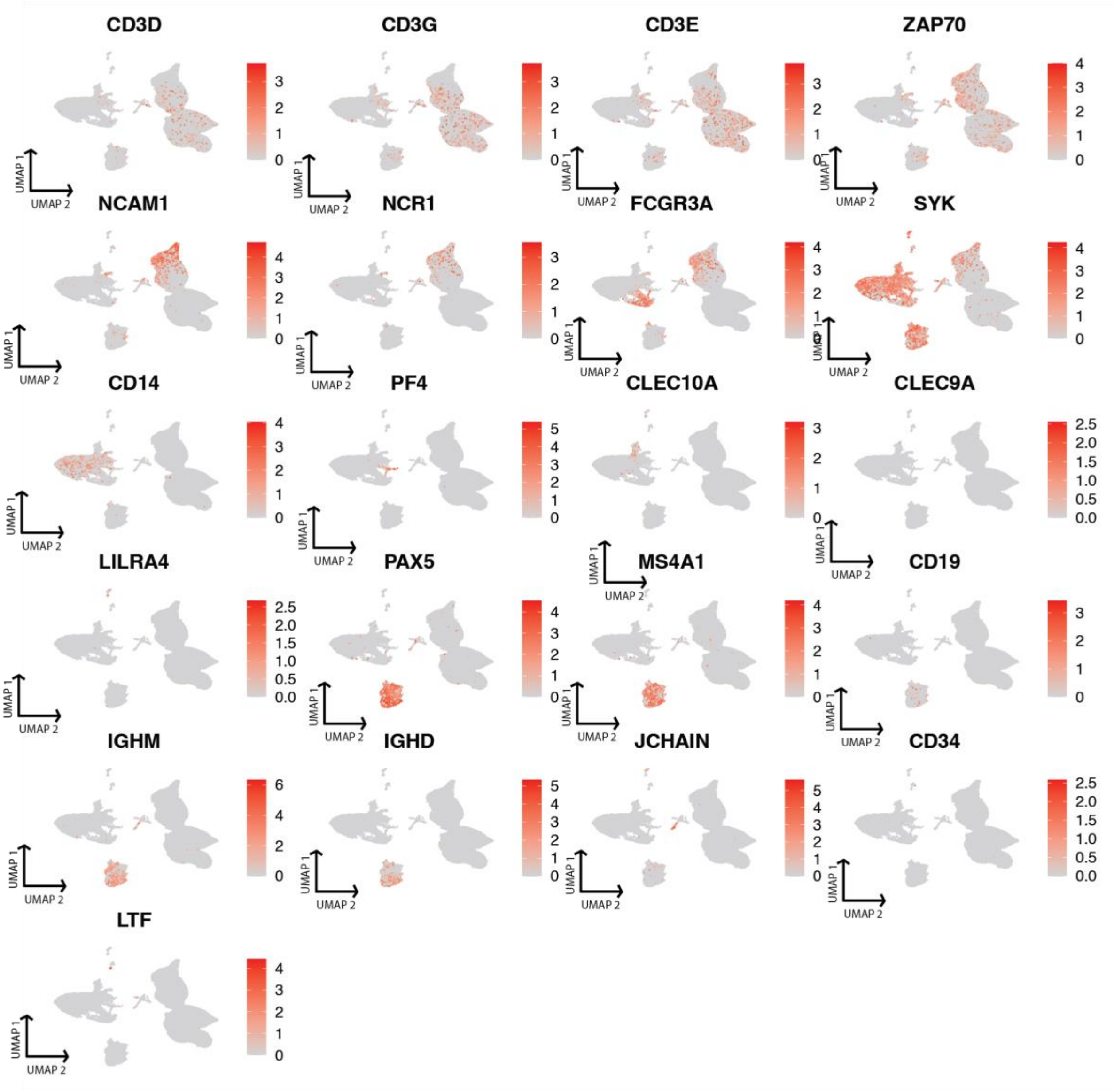


**Supplemental Figure 2: Feature plots of the expression of each broad marker genes used for the first level of cell type annotation.** Each subplot depicts the expression of a marker gene graphically on a UMAP consisting of all PBMCs from all 15 participants.

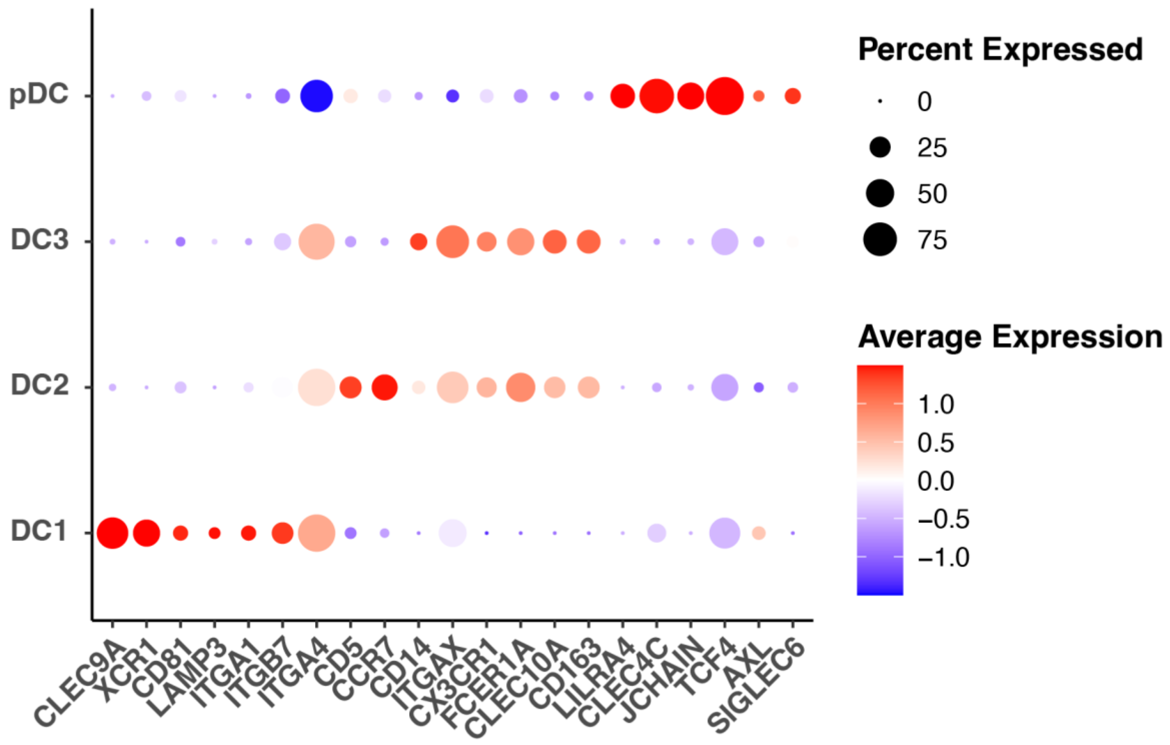


**Supplemental Figure 3: Dendritic Cell (DC) clusters**. Dot plot showing marker genes used in the second level of annotation to DC cell subpopulations including DC type 1 (DC1), DC type 2 (DC2), DC type 3 (DC3), and plasmacytoid dendritic cell (pDC), with color bar indicating relative z-scored expression of each marker gene and the dot size indicating the percentage of cells in that cluster expressing that marker gene.

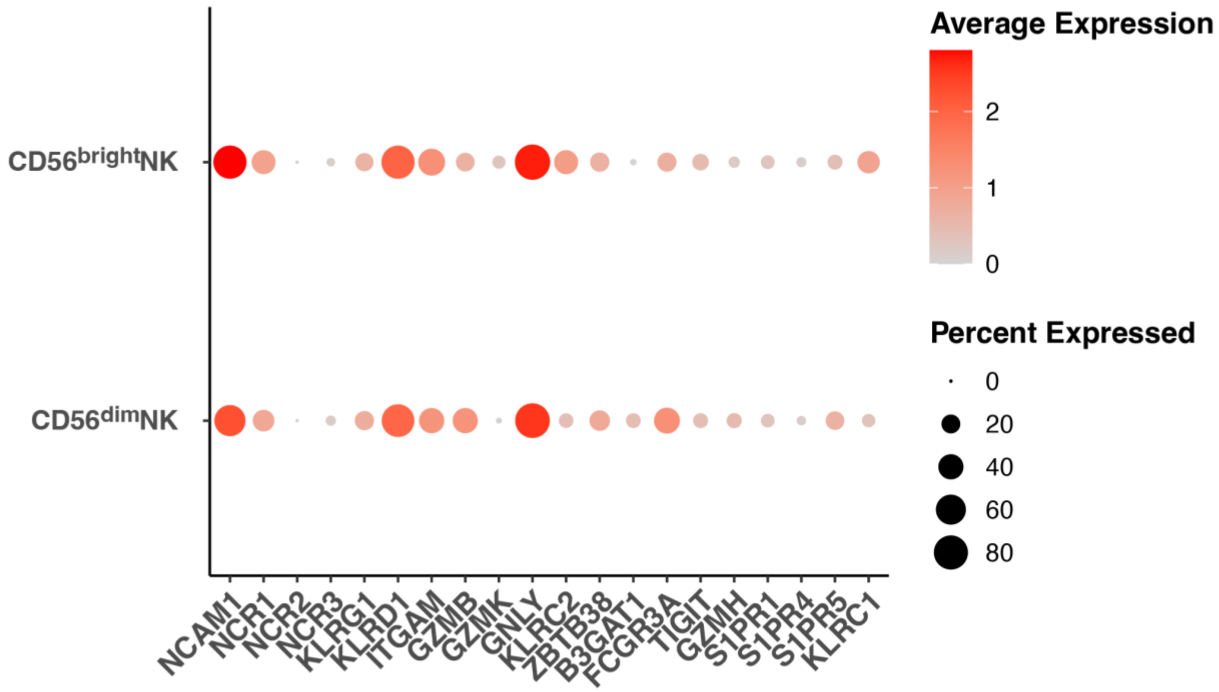


**Supplemental Figure 4: Natural Killer (NK) cell clusters.** Dot plot showing marker genes used in the second level of annotation to NK subpopulations including NK CD56$^{bright}$ and NK CD56$^{dim}$ subpopulations, with color bar indicating average expression of each marker gene.

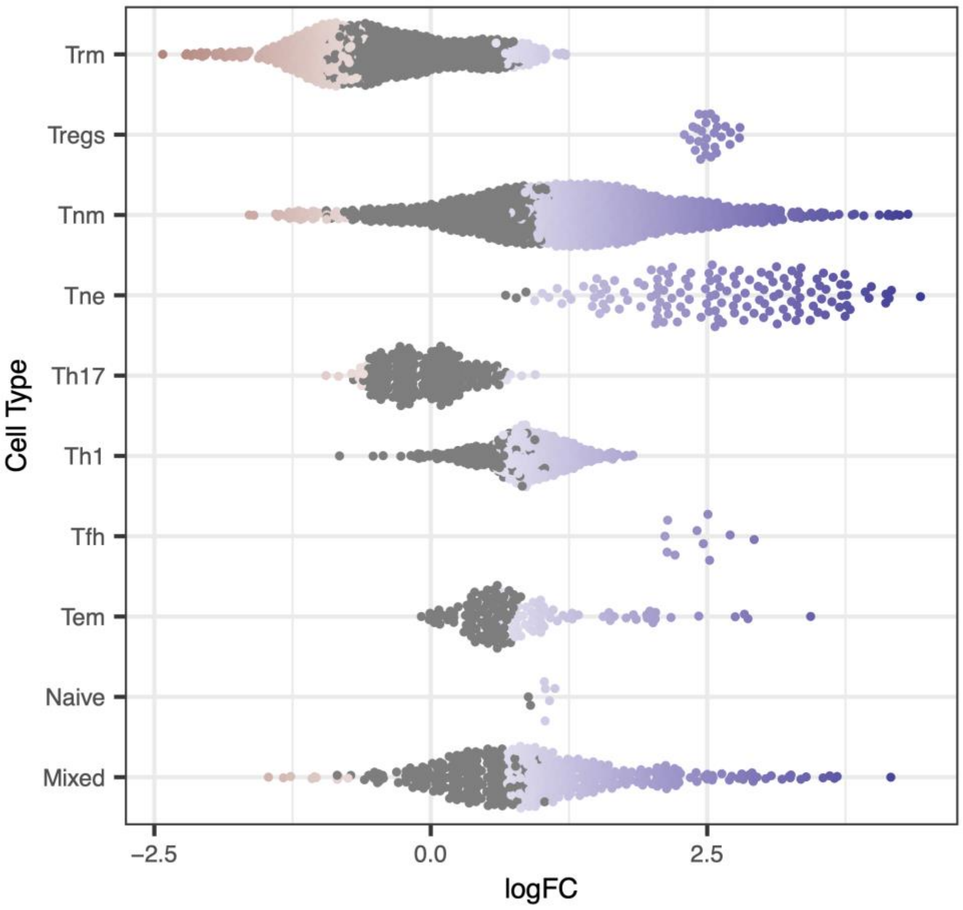


**Supplemental Figure 5: Kornberg miloR results.** Beeswarm plot of differential abundance analysis results using Milo on Kornberg et al. (2023) single-cell RNA-seq. Each dot represents a neighborhood, and the log-fold change represents the decrease or increase in a neighborhood in Celiac disease compared to healthy controls.

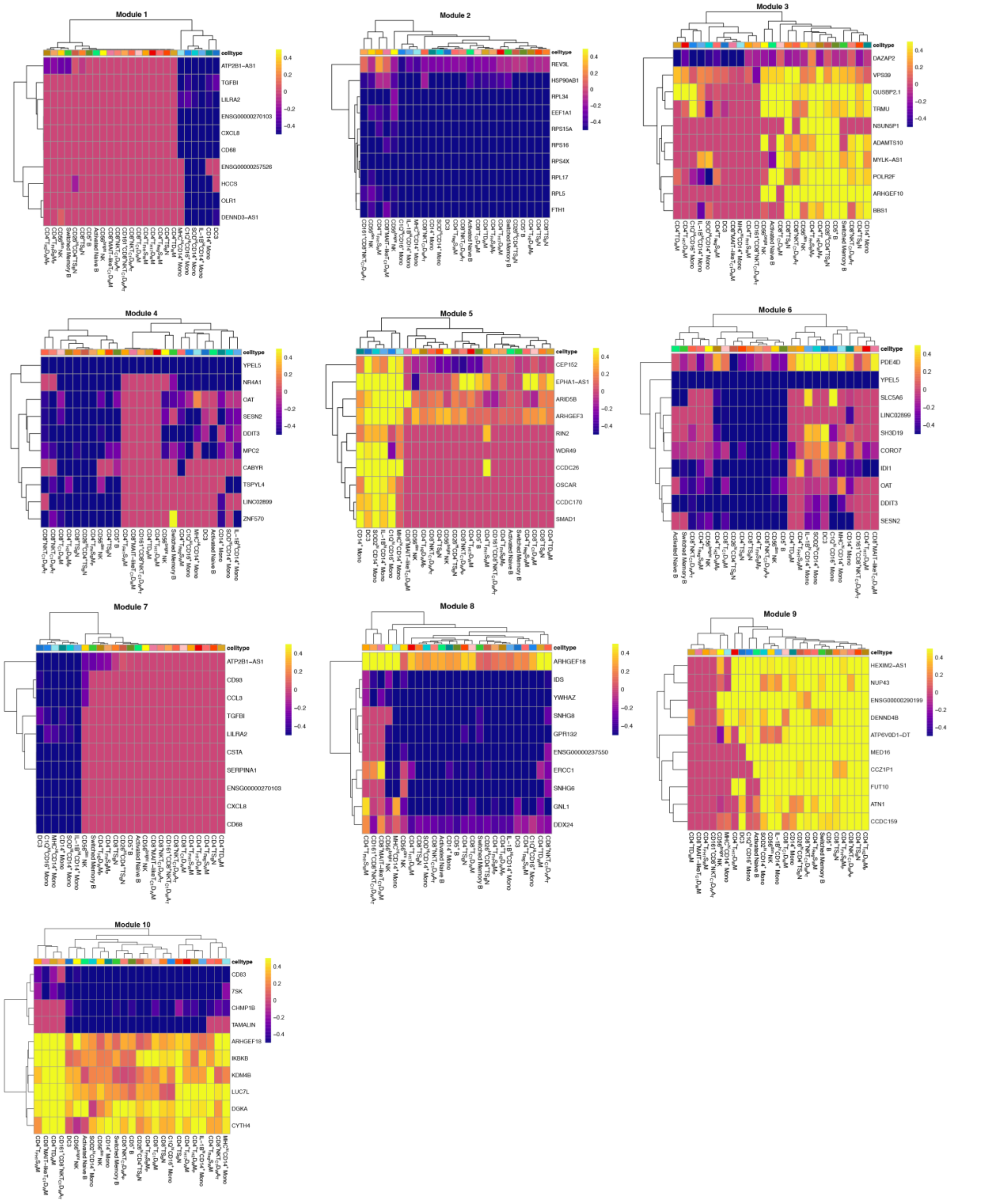


**Supplemental Figure 6: K-means Clustering of Differentially Expressed Genes into Transcriptional Modules.** Heatmaps for the top 10 genes in each module, colored by the relative upregulation or downregulation in AGA+ SRD as compared to AGA- participants (AGA- SRD and AGA- HC). Warmer colors represent modules upregulated in that cell type in AGA+ SRD compared to AGA- participants. The columns are annotated with immune cell types. Module 1 corresponds to genes in the regulation of translation pathway, module 2 corresponds to genes in cytoplasmic translation pathway, module 3 corresponds to genes in the cellular response to chemical stress pathway, module 4 corresponds to genes in the purine ribonucleotide catabolic process pathway, module 5 corresponds to genes in the arginine catabolic process pathway, module 6 corresponds to genes in the growth hormone receptor signaling pathway via JAK-STAT pathway, module 7 corresponds to genes in the protein acylation pathway, module 8 corresponds to genes in the regeneration pathway, module 9 corresponds to genes in the response to lipopolysaccharide pathway, and module 10 corresponds to genes in the positive regulation of type 1 interferon-mediated signaling pathway.

### Supplemental Tables

#### Table 1: Demographic and Clinical Characteristics of Participants

| Participant ID | Age (years ) | Sex | Race | Primary Psychiatric Diagnosis | Secondary Psychiatric Diagnoses | Immunological Medical History | AGA-IgG (U) | AGA Status | Group |
|---|---|---|---|---|---|---|---|---|---|
| M021407 | 53 | M | White | Schizophrenia | None | None | 124.40 | Positive | AGA+ SRD |
| M031465 | 53 | M | White | Schizophrenia | None | None | 0.95 | Negative | AGA- SRD |
| M064023 | 39 | M | Mixed | Schizophrenia | None | None | 6.40 | Negative | AGA- SRD |
| M070176 | 38 | M | White | Schizophrenia | None | None | 129.50 | Positive | AGA+ SRD |

| | | | | | | | | | |
|---|---|---|---|---|---|---|---|---|---|
| M078671 | 37 | M | White | Schizophrenia | None | None | 23.64 | Positive | AGA+ SRD |
| M085678 | 46 | M | Black or African American | Schizophrenia | None | None | 31.87 | Positive | AGA+ SRD |
| M085836 | 39 | M | White | Schizophrenia | None | None | 32.10 | Positive | AGA+ SRD |
| M089673 | 46 | F | Black or African American | Schizoaffective disorder, depressed type | Post-Traumatic Stress Disorder (PTSD) | None | 84.52 | Positive | AGA+ SRD |
| M093330 | 37 | F | Black or African American | Schizophrenia | Alcohol Use Disorder (AUD) | History of HIV Positive on Anti-Retroviral Therapy | 4.82 | Negative | AGA- SRD |
| M093701 | 35 | F | Hispanic | Schizoaffective disorder, bipolar type | None | N/A | 22.97 | Positive | AGA+ SRD |
| M093914 | 36 | F | White | N/A | N/A | N/A | 12.93 | Negative | HC |
| M099090 | 46 | M | Asian | N/A | N/A | N/A | 19.33 | Negative | HC |
| M099325 | 25 | M | White | N/A | N/A | N/A | 6.33 | Negative | HC |

| | | | | | | | | | |
|---|---|---|---|---|---|---|---|---|---|
| M099786 | 35 | M | White | N/A | N/A | N/A | 9.09 | Negative | HC |
| M100776 | 53 | F | White | N/A | N/A | N/A | 8.55 | Negative | HC |

**Supplemental Table 1: Demographic and clinical characteristics of participants.** Participants in order by increasing ID number. Abbreviations: Anti-Gliadin Antibody Immunoglobulin G (AGA IgG), Schizophrenia Related Disorders (SRD), Healthy Controls (HC)

| Variable | AGA+ SRD (n=7) | AGA- SRD (n=3) | HC (n=5) | Statistics |
|---|---|---|---|---|
| Age (years, mean ± SD) | 42 ± 6.48 | 43 ± 8.72 | 39 ± 10.79 | F(2,12)= 0.27; p= 0.77 (ns) |
| Sex | Male (n=5), Female (n=2) | Male (n=3), Female (n=1) | Male (n=3), Female (n=2) | p= 1.01 (ns) |
| Race | White (n=4) Black or African American (n=2), Hispanic (n=1) | White (n=1), Black or African American (n=1), Mixed (n=1) | White (n=4), Asian (n=1) | p= 0.38 (ns) |
| AGA-IgG (U) | 64.14 ± 47.82 | 4.06 ± 2.80 | 11.25 ± 5.11 | F(2,12)= 5.04; p= 0.026* |

**Supplemental Table 2: Group Comparisons of Demographic Variables.** One-way ANOVA was used to compare age or AGA-IgG across AGA+ SRD, AGA- SRD, and HC. Fisher's Exact used to compare Sex and Race across AGA+ SRD, AGA- SRD, and HC. Abbreviations: ns= not significant. *Indicates p<0.05.

| Cluster Label | Positive Markers | Negative Markers |
|---|---|---|
| Dendritic Cell type 1 (DC1) | *CLEC9A, XCR1, ITGA1, ITGA4, ITGB7, LAMP3, CD81, CD5, CD163, CCR7, ITGAX, CX3CR1, AXL* | *CD5, CLEC10A, LILRA4, CLEC4C, JCIAN, TCF4, CD14, SIGLEC6, CCR7, FCER1A* |

| | | |
|---|---|---|
| Dendritic Cell type 2 (DC2) | *CLEC10A, CD5, CCR7, CD163, ITGAX, CX3CR1, ITGA4, FCER1A* | *CLEC9A, XCR1, CD81, LAMP3, LILRA4, CLEC4C, JCHAIN, TCF4, CD14, AXL, SIGLEC6, ITGA1, ITGB7, AXL, SIGLEC6* |
| Dendritic Cell type 3 (DC3) | *CLEC10A, CD14, CD163, ITGAX, ITGA4, FCER1A* | *CD5, CCR7, CLEC9A, XCR1, CD81, LAMP3, LILRA4, CLEC4C, JCHAIN, TCF4, ITGA1, ITGB7, AXL, SIGLEC6* |
| Plasmacytoid Dendritic Cell (pDC) | *LILRA4, CLEC4C, JCHAIN, TCF4, AXL, SIGLEC6* | *CLEC9A, XCR1, CD81, LAMP3, ITGA1, ITGB7, ITGA4, CD5, CCR7, CD14, ITGAX, CX3CR1, FCER1A, CLEC10A, CD163* |

**Supplemental Table 3: Dendritic Cell (DC) annotation marker genes (second level).**

| **Cluster Label** | **Positive Markers** | **Negative Markers** |
|---|---|---|
| Activated Naive B cells | *MS4A1, TCL1A, YBX3, FCER2, IL4R, CD38, CXCR5, CD69, CD83, IGHM, IGHD, CD19, CR2, CD24, JUN* | *CD27, AIM2, TNFRSR13B, IGHG1, IGHG2, IGHG3, IGHG4, IGHA1, IGHA2, FAS, CD5, LEF1, TBX21, SPN, ITGAX, ZEB2, MEF2B, CD86, BCL6, LMO2, FOS, JUNB* |
| Switched Memory B cells | *MS4A1, CXCR5, CD19, CR2, CD24, CD27, AIM2, TNFRSR13B, IGHG1, IGHG2, IGHG3, IGHG4, IGHA1, IGHA2, FAS* | *TCL1A, YBX3, FCER2, IL4R, CD38, CD69, CD83, IGHM, IGHD, CD5, LEF1, TBX21, SPN, ITGAX, ZEB2, MEF2B, CD86, BCL6, LMO2, FOS, JUN, JUNB* |
| CD5+ B cells | *CD69, CD27, FAS, CD5, LEF1, TBX21, SPN* | *MS4A1, TCL1A, YBX3, IL4R, CD38, CD83, IGHM, IGHD, CD19, CR2, CD24, AIM2, TNFRSR13B, IGHG1, IGHG2, IGHG3, IGHG4, IGHA1, IGHA2, ITGAX, ZEB2, MEF2B, CD86, BCL6, LMO2, FOS, JUN, JUNB* |

| | | |
|---|---|---|
| Atypical Memory B cells | *TGAX, ZEB2, CD86, BCL6, TBX21, SPN, ITGAX, ZEB2, MEF2B, CD86, BCL6, LMO2, FOS, JUN, JUNB* | *MS4A1, TCL1A, FCER1, IL4R, CD38, CXCR5, CD69, CD83, IGHM, IGHD,*<br>*CD19, CR2, CD24, CD27, AIM2, TNFRSF13B, IGHG1, IGHG2, IGHG3, IGHG4, IGHA1, IGHA2, FAS* |
| Plasmablasts | *CD27, IGHM, TNFRSF13B, JCHAIN, IGHG1, IGHG2, IGHG3, IGHG4, IGHA1, FAS, TBX21, SPN,*<br>*JUN, TXNDC5, JCHAIN, MZB1, XBP1, PRDM1, CD38* | *MS4A1, CD19, CD24* |

**Supplemental Table 4: B cell annotation marker genes (second level)**

| Cluster Label | Higher relative expression | Lower relative expression |
|---|---|---|
| CD56-dim NK cells | *FCGR3A, GZMB, CX3CR1, LILRB1, B3GAT1* | GZMK, KLRC1, KLRC2, IL7R, SELL |
| CD56-bright NK cells | *NCAM1, GZMK, KLRC1, KLRC2, IL7R, SELL* | *FCGR3A, GZMB, CX3CR1, B3GAT1, LILRB1* |

**Supplemental Table 5: NK cell annotation marker genes (second level)**

| Cluster Label | Positive/Brighter Expression Markers | Negative/Lower Expression Markers |
|---|---|---|
| CD14+ Monocytes | *CD14, S100A8, S100A9, SOD2, MGST1, HLA-A, IL1B, CCL3* | *FCGR3A, S100A12, NAMPT, CST3, HLA-C, HLA-DRA, HLA-DRB1, HLA-DQA1, HLA-DQB1, HLA-DPA1, HLA-* |

| | | |
|---|---|---|
| | | *DPB1, IRF8, ICAM1, JUNB, EGR3, CCR2, ISG15, IFI44L, MX1, MX2, MS4A7, CX3CR1, STAT1, C1QA, C1QB, CD68* |
| MHC-Hi CD14+ Monocytes | *CD14, S100A8, S100A9, S100A12, MGST1, CST3, HLA-A, HLA-B, HLA-C, HLA-DRA, HLADR-B1, HLA-DQA1, HLA-DQB1, HLA-DPA1, IL1B, IRF8, ICAM1, JUNB, CCL3, CCR2, CD68* | *FCGR3A, SOD2, NAMPT, EGR3, ISG15, IFI44L, MX1, MX2, MS4A7, CX3CR1, C1QA, C1QB* |
| SOD2-Hi CD14+ Monocytes | *CD14, S100A8, S100A9, S100A12, SOD2, MGST1, HLA-A, IL1B, IRF8, JUNB, ICAM1, JUNB, EGR3, CCR2, CD68* | *FCGR3A, CST3, HLA-A, HLA-C, HLA-DRA, HLADR-B1, HLA-DQA1, HLA-DQB1, HLA-DPA1, HLA-DPB1, IGS15, IFI44L, MX1, MX2, MS4A7, CX3CR1, C1QA, C1QB* |
| IL-1B-Hi CD14+ Monocytes | *CD14, S100A8, S100A9, SOD2, MGST1, HLA-A, HLA-B, HLA-C, HLA-DRA, HLADR-B1, HLA-DQA1, HLA-DQB1, HLA-DPA1, IL1B, CST3, IRF8, JUNB, ICAM1, JUNB, EGR3, CCL3, CCR2, CD68* | *FCGR3A, S100A12, FCGR3A, CST3, IGS15, IFI44L, MX1, MX2, MS4A7, CX3CR1, C1QA, C1QB* |
| ISG+ CD14+ Monocytes | *CD14, S100A8, S100A9, S100A12, SOD2, NAMPT, CST3, HLA-A, HLC-C, HLA-DRA, HLA-DRB1, HLA-DQA1, HLA-DQB1, HLA-DPA1, HLA-DPB1, IL1B, IRF8, ICAM1, CCL3, CCR2, ISG15, IFI44L, MX1, MX2, CD68* | *FCG3RA, MGST1, HLA-B, JUNB, ERG3, MS4A7, CX3CR1, C1QA, C1QB, C1QC* |
| CD16+ Monocytes | *FCGR3A, MS4A7, CX3CR1, C1QA, CQ1B* | *CD14, S100A8, S100A9, S100A12, SOD2, MGST1, NAMPT, CST3, MGST1, HLA-A, HLA-B, HLA-C, HLA-DRA, HLADR-B1, HLA-DQA1, HLA-DQB1, HLA-DPA1, IL1B,* |

| | | |
|---|---|---|
| | | *IRF8, ICAM1, JUNB, EGR3, CCL3, CCR2, ISG15, IFI44L, MX1, MX2, SIGLEC1, STAT1, CD68* |
| C1Q-Hi CD16+ Monocytes | *FCGR3A, NAMPT, CST3, HLA-C, HLA-DRA, HLA-DQA1, HLA-DPA1, HLA-DPB1, JUNB, MS4A7, CX3CR1, C1QA, CQ1B, CD68* | *CD14, S100A8, S100A9, S100A12, SOD2, MGST1, HLA-A, HLA-B, HLA-DRB1, HLA-DQB1, IL1B, IRF8, ICAM1, EGR3, CCL3, CCR2, ISG15, IFI44L, MX1, MX2, SIGLEC1, STAT1* |

**Supplemental Table 6: Monocyte annotation marker genes (second level)**

**Supplemental Table 7: T cell annotation marker genes (second level)**

| **Cluster Label** | **Higher relative expression** | **Lower relative expression** |
|---|---|---|
| $CD26^{Hi}$ $CD4^{+}$ $TS_{B}N$ | *CD4, IL6ST, FOXO1, TCF7, IL7R, CD28, LEF1, DPP4, CCR7, SELL, ICOS* | *CD8A, CD8B, KLRB1, ZBTB16, SLC4A10, FAS, CXCR4, CD44, IL2RA, IL2RB, KLF2, S1PR1, CD69, ITGA1, ITGA4, ITGAE, ITGB1, ITGB7, ABCB1, CXCR3, GATA3, CCR4, PTGDR2, BCL6, MAF, CXCR5, IKZF2, FOXP3, CCR8, CTLA4, TIGIT, HAVCR2, TNFRSF1B, CCR10, AHR, CCR6, RORC, IL23R, KIT, IL10RA, IL18R1, IL18RAP, NCAM1, B3GAT1, CD160, EOMES, TBX21, IFNG, CCR5, CXRCR1, FCGR3A, FCER1G, KLRK1, GZMK, KLRC1, KLRC2, PRF1, KLRG1, GZMB, GNLY, FGFBP2, ZEB2, PRDM1, LAG3, PDCD1, FASLG, FGL2* |
| $CD8^{+}$ $TS_{B}N$ | *CD8A, CD8B, IL6ST, FOXO1, TCF7, CD27, IL7R, LEF1, CCR7, SELL, ABCB1, KLRK1* | *CD4, KLRB1, ZBTB16, SLC4A10, CD28, FAS, CXCR4, CD44, IL2RA, IL2RB, KLF2, DPP4, S1PR1, CD69, ITGA1, ITGA4, ITGAE, ITGB1, ITGB7, CXCR3,* |

| | | |
|---|---|---|
| | | *GATA3, CCR4, PTGDR2, BCL6, MAF, ICOS, CXCR5, IKZF2, FOXP3, CCR8, CTLA4, TIGIT, HAVCR2, TNFRSF1B, CCR10, AHR, CCR6, RORC, IL23R, KIT, IL10RA, IL18R1, IL18RAP, NCAM1, B3GAT1, CD160, EOMES, TBX21, IFNG, CCR5, CXRCR1, FCGR3A, FCER1G, GZMK, KLRC1, KLRC2, PRF1, KLRG1, GZMB, GNLY, FGFBP2, ZEB2, PRDM1, LAG3, PDCD1, FASLG, FGL2* |
| CD4$^+$ TS$_B$N | *CD4, IL6ST, FOXO1, TCF7, CD27, IL7R, CD28, LEF1, DPP4, CCR7, SELL, ICOS* | *CD8A, CD8B, KLRB1, ZBTB16, SLC4A10, FAS, CXCR4, CD44, IL2RA, IL2RB, S1PR1, CD69, ITGA1, ITGA4, ITGAE, ITGB1, ITGB7, ABCB1, CXCR3, GATA3, CCR4, PTGDR2, BCL6, MAF, CXCR5, IKZF2, FOXP3, CCR8, CTLA4, TIGIT, HAVCR2, TNFRSF1B, CCR10, AHR, CCR6, RORC, IL23R, KIT, IL10RA, IL18R1, IL18RAP, NCAM1, B3GAT1, CD160, EOMES, TBX21, IFNG, CCR5, CXRCR1, FCGR3A, FCER1G, KLRK1, GZMK, KLRC1, KLRC2, PRF1, KLRG1, GZMB, GNLY, FGFBP2, ZEB2, PRDM1, LAG3, PDCD1, FASLG, FGL2* |
| CD4$^+$ T$_{H2}$D$_B$M$_P$ | *CD4, ZBTB16, IL6ST, TCF7, IL7R, CD28, LEF1, FAS, CD44, IL2RA, KLF2, SELL, S1PR1, CD69, ITGA4, ITGB1, GATA3, CCR4, PTGDR2, MAF,* | *CD8A, CD8B, KLRB1, SLC4A10, FOXO1, CD27, CXCR4, IL2RB, DPP4, CCR7, ITGA4, ITGB7, ABCB1, CXCR3, BCL6, CXCR5, IKZF, FOXP3, CTLA4, TIGIT, HAVCR2, TNFRSF1B, CCR6, RORC, IL23R, KIT, IL18R1, IL18RAP, NCAM1, B3GAT1,* |

| | | |
|---|---|---|
| | *ICOS CCR8, CCR10, AHR, IL10RA, PRDM1* | *CD160, EOMES, TBX21, IFNG, CCR5, CXRCR1, FCGR3A, FCER1G, KLRK1, GZMK, KLRC1, KLRC2, PRF1, KLRG1, GZMB, GNLY, FGFBP2, ZEB2, LAG3, PDCD1, FASLG, FGL2* |
| $CD4^{+}T_{FH}\ S_{B}M_{P}$ | *CD4, IL6ST, FOXO1, TCF7, IL7R, CD28, LEF1, FAS, CD44, IL2RA, DPP4, CCR7, SELL, S1PR1, CD69, ITGA4, ITGB1, CCR4, MAF, ICOS, CXCR5, AHR* | *CD8A, CD8B, KLRB1, ZBTB16, SLC4A10, CD27, CXCR4, IL2RB, KLF2, ITGA1, ITGAE, ITGB7, ABCB1, CXCR3, GATA3, BCL6, IKZF2, FOXP3, CCR8, CTLA4, TIGIT, HAVCR2, TNFRSF1B, CCR10, CCR6, RORC, IL23R, KIT, IL10RA, IL18R1, IL18RAP, NCAM1, B3GAT1, CD160, EOMES, TBX21, IFNG, CCR5, CXRCR1, FCGR3A, FCER1G, KLRK1, GZMK, KLRC1, KLRC2, PRF1, KLRG1, GZMB, GNLY, FGFBP2, ZEB2, PRDM1, LAG3, PDCD1, FASLG, FGL2* |
| $CD8^{+}\ T_{C1}\ D_{W}\ M$ | *CD8A, CD8B, ZBTB16, IL7R, FAS, CXCR4, CD44, IL2RB, DPP4, CD69, ITGA1, ITGA4, ITGAE, ABCB1, CXCR3, BCL6, TIGIT, AHR, IL10RA, IL18R1, IL18RAP, CD160, EOMES, TBX21, IFNG, CCR5, KLRK1, GZMK, KLRC1, KLRC2, KLRG1, GNLY, ZEB2, PRDM1, LAG3, PDCD1* | *CD4, KLRB1, SLC4A10, IL6ST, FOXO1, TCF7, CD27, CD28, LEF1, IL2RA, KLF2, CCR7, SELL, S1PR1, ITGB1, ITGB7, GATA3, CCR4, PTGDR2, MAF, ICOS, CXCR5, IKZF2, FOXP3, CCR8, CTLA4, HAVCR2, TNFRSF1B, CCR10, CCR6, RORC, IL23R, KIT, NCAM1, B3GAT1, CD160, EOMES, TBX21, IFNG, CCR5, CX3CR1, KLRK1, GZMK, KLRC1, KLRC2, KLRG1, GNLY, ZEB2, PRDM1, LAG3, PDCD1* |
| $CD8^{+}NKT_{C1}\ D_{W}\ A_{t}$ | *CD8A, CD8B, IL2RB, KLF2, S1PR1, ITGA4, ITGB1, ITGB7, ABCB1,* | *CD4, KLRB1, ZBTB16, SLC4A10, IL6ST, FOXO1, TCF7, CD27, IL7R, CD28, LEF1, FAS, CXCR4,* |

| | | |
|---|---|---|
| | *CXCR3, IKZF2, TIGIT, TNFRSF1B, IL10RA, IL18R1, IL18RAP, NCAM1, B3GAT1, CD160, EOMES, TBX21, IFNG, CX3CR1, FCGR3A, FCER1G, KLRK1, KLRC1, KLRC2, PRF1, KLRG1, GZMB, GNLY, FGFBP2, ZEB2, PRDM1, LAG3, PDCD1, FASLG, FGL2* | *CD44, IL2RA, DPP4, CCR7, SELL, ITGA1, ITGAE, GATA3, CCR4, PTGDR2, BCL6, MAF, ICOS, CXCR5, FOXP3, CCR8, CTLA4, HAVCR2, CCR10, AHR, CCR6, RORC, IL23R, KIT, CCR5, GZMK* |
| $CD4^+$ $T_{H17}$ $D_WM$ | *CD4, KLRB1, ZBTB16, SLC4A10, FOXO1, IL7R, CD28, CD44, IL2RA, KLF2, DPP4, CD69, ITGA1, ITGA4, ITGAE, ITGB1, ITGB7, ABCB1, GATA3, MAF, AHR, CCR6, RORC, IL23R, KIT, IL10RA, IL18R1, PRDM1* | *CD8A, CD8B, IL6ST, TCF7, CD27, LEF1, FAS, CXCR4, IL2RB, CCR7, SELL, S1PR1, CXCR3, CCR4, PTGDR2, BCL6, ICOS, CXCR5, IKZF2, FOXP3, CCR8, CTLA4, TIGIT, HAVCR2, TNFRSF1B, CCR10, IL18RAP, NCAM1, B3GAT1, CD160, EOMES, TBX21, IFNG, CCR5, CXRCR1, FCGR3A, FCER1G, KLRK1, GZMK, KLRC1, KLRC2, PRF1, KLRG1, GZMB, GNLY, FGFBP2, ZEB2, PRDM1, LAG3, PDCD1, FASLG, FGL2* |
| $CD8^+$MAIT-like-$T_{C17}D_WM$ | *CD8A, KLRB1, ZBTB16, SLC4A10, IL7R, CXCR4, CD44, IL2RB, DPP4, S1PR1, CD69, ITGA1, ITGAE, ITGB7, ABCB1, GATA3, BCL6, MAF, IKZF2, CCR6, RORC, IL23R, KIT, IL18R1, IL18RAP, NCAM1, EOMES, TBX21, CCR5, KLRK1, GZMK, PRF1, KLRG1, PRDM1, LAG3, FASLG* | *CD4, CD8B, IL6ST, FOXO1, TCF7, CD27, CD28, LEF1, FAS, IL2RA, KLF2, CCR7, SELL, ITGA4, ITGB1, CXCR3, CCR4, PTGDR2, ICOS, CXCR5, FOXP3, CCR8, CTLA4, TIGIT, HAVCR2, TNFRSF1B, CCR10, AHR, IL10RA, B3GAT1, CD160, IFNG, CX3CR1, FCGR3A, FCER1G, KLRC2, GZMB, FGFBP2, ZEB2, FGL2* |

| | | |
|---|---|---|
| $CD4^{+}$ $TD_{W}M$ | *CD4, IL6ST, TCF7, IL7R, CD28, FAS, CXCR4, CD44, KLF2, DPP4, SELL, CD69, S1PR1, ITGA1, ITGA4, ITGAE, ITGB1, ITGB7, CXCR3, GATA3, CCR4, PTGDR2, BCL6, MAF, ICOS, CXCR5, CCR10, AHR, CCR6, IL10RA, PRDM1* | *CD8A, CD8B, KLRB1, ZBTB16, SLC4A10, FOXO1, CD27, IL7R, CD28, FAS, CXCR4, IL2RB, CCR7, S1PR1, ABCB1, IKZF2, FOXP3, CCR8, CTLA4, TIGIT, HAVCR2, TNFRSF1B, RORC, IL23R, KIT, IL18R1, IL18RAP, NCAM1, B3GAT1, CD160, EOMES, TBX21, IFNG, CCR5, CXRCR1, FCGR3A, FCER1G, KLRK1, GZMK, KLRC1, KLRC2, PRF1, KLRG1, GZMB, GNLY, FGFBP2, ZEB2, LAG3, PDCD1, FASLG, FGL2* |
| $CD161^{+}CD8^{+}NKT_{C1}D_{W}A_{t}$ | *CD8A, CD8B, KLRB1, ZBTB16, IL2RB, KLF2, S1PR1, ITGAE, ITGB7, ABCB1, CXCR3, PTGDR2, BCL6, MAF, IKZF2, TIGIT, RNFRSF1B, IL18R1, IL18RAP, NCAM1, B3GAT1, CD160, EOMES, TBX21, IFNG, CCR5, CX3CR1, FCGR3A, FCER1G, KLRK1, GZMK, KLRC1, KLRC2, PRF1, KLRG1, GZMB, GNLY, FGFBP2, ZEB2, PRDM1, LAG3, PDCD1, FASLG, FGL2* | *CD4, SLC4A10, IL6ST, FOXO1, TCF7, CD27, IL7R, CD28, LEF1, FAS, CXCR4, CD44, IL2RA, DPP4, CCR7, SELL, CD69, ITGA1, ITGA4, ITGB1, GATA3, CCR4, ICOS, FOXP3, CCR8, CTLA4, HAVCR2, CCR10, AHR, CCR6, RORC, IL23R, KIT, IL10RA* |
| $CD4^{+}$ $T_{reg}$ $S_{W}$ M | *CD4, IL6ST, FOXO1, CD27, CD28, LEF1, FAS, CD44, IL2RA, IL2RB, SELL, ITGB1, GATA3, CCR4, MAF, ICOS,* | *CD8A, CD8B, KLRB1, ZBTB16, SLC4A10, TCF7, IL7R, CXCR4, KLF2, DPP4, CCR7, S1PR1, CD69, ITGA1, ITGA4, ITGAE, ITGB7, ABCB1, CXCR3,* |

| | | |
|---|---|---|
| | *IKZF2, FOXP3, CCR8, CTLA4, TIGIT, TNFRSF1B, CCR10, AHR, CCR6, IL10RA, PRDM1* | *PTGDR2, BCL6, CXCR5, HAVCR2, RORC, IL23R, KIT, IL18R1, IL18RAP, NCAM1, B3GAT1, CD160, EOMES, TBX21, IFNG, CCR5, CXRCR1, FCGR3A, FCER1G, KLRK1, GZMK, KLRC1, KLRC2, PRF1, KLRG1, GZMB, GNLY, FGFBP2, ZEB2, LAG3, PDCD1, FASLG, FGL2* |
| $CD8^{+}NKT_{C1}$ $D_{W}$ $A_{P}$ | *CD8A, CD8B, ZBTB16, IL6ST, LEF1, IL2RB, KLF2, CCR7, ITGB1, ABCB1, IKZF2, HAVCR2, TNFRSF1B, IL18R1, IL18RAP, NCAM1, B3GAT1, CD160, EOMES, TBX21, IFNG, CX3CR1, FCGR3A, FCER1G, KLRK1, KLRC1, KLRC2, PRF1, KLRG1, GZMB, GNLY, FGFBP2, ZEB2, FASLG, FGL2* | *CD4, KLRB1, SLC4A10, FOXO1, TCF7, CD27, IL7R, CD28, FAS, CXCR4, CD44, IL2RA, DPP4, SELL, S1PR1, CD69, ITGA1, ITGA4, ITGAE, ITGB7, CXCR3, GATA3, CCR4, PTGDR2, BCL6, MAF, ICOS, CXCR5, FOXP3, CCR8, CTLA4, TIGIT, CCR10, AHR, CCR6, RORC, IL23R, KIT, IL10RA, CCR5, GZMK, PRDM1, PDCD1* |
| $CD4^{-}CD8^{dim}$ $T_{Reg}$ $S_{W}M_{P}$ | *CD8B, FOXO1, TCF7, CD27, CD28, LEF1, FAS, CD44, CCR7, SELL, S1PR1, CD69, ITGA4, ITGAE, ITGB1, ITGB7, CXCR3, MAF, CXCR5, CTLA4, TIGIT, HAVCR2, TNFRSF1B, B3GAT1, EOMES, CCR5, KLRK1, GZMK, LAG3, PDCD1, CD38, MKI67* | *CD4, CD8A, KLRB1, ZBTB16, SLC4A10, TCF7, IL7R, CXCR4, KLF2, DPP4, ITGA1, ABCB1, GATA3, CCR4, PTGDR2, BCL6, IKZF2, FOXP3, CCR8, CCR10, AHR, CCR6, RORC, IL23R, KIT, IL10RA, IL18R1, IL18RAP, NCAM1, CD160, TBX21, IFNG, CX3CR1, FCGR3A, FCER1G, KLRC1, KLRC2, PRF1, KLRG1, GZMB, GNLY, FGFBP2, ZEB2, PRDM1* |
| $CD4^{+}T_{FH1}S_{B}M$ | *CD4, CD8A, IL6ST, LEF1, CD44, KLF2, DPP4, CCR7, SELL, ITGAE,* | *CD8B, FOXO1, TCF7, CD27, IL7R, CD28, FAS, CXCR4, IL2RA, IL2RB, DPP4, S1PR1,* |

| | | |
|---|---|---|
| | *CXCR3, BCL6, HAVCR2, TNFRSF1B, AHR, NCAM1, CX3CR1, FCGR3A, FCER1G, GNLY, ZEB2, FGL2* | *CD69, ITGA1, ITGA4, ITGB1, ITGB7, ABCB1, GATA3, CCR4, PTGDR2, MAF, ICOS, CXCR5, IKZF2, FOXP3, CCR8, CTLA4, TIGIT, HAVCR2, CCR10, CCR6, RORC, IL23R, KIT, IL10A, IL18R1, IL18RAP, B3GAT1, CD160 EOMES, TBX21, IFNG, CCR5, KLRK1, GZMK, KLRC1, KLRC2, PRF1, KLRG1, GZMB, FGFBP2, PRDM1, LAG3, PDCD1, FASLG* |
| Ki67$^+$ CD8$^+$NKT$_{C1}$D$_W$A$_t$ | *CD8A, KLRB1, ZBTB16, IL2RB, SELL, ITGA4, ITGAE, ITGB1, ITGB7, CXCR3, CCR4, PTDGR2, ICOS, IKZF2, CCR8, CTLA4, TIGIT, HAVCR2, TNFRSF1B, CCR10, KIT, IL18RAP, NCAM1, B3GAT1, CD160, EOMES, TBX21, IFNG, CCR5, CX3CR1, FCGR3A, FCER1G, GZMK, KLRK1, KLRC1, KLRC2, PRF1, GZMB, GNLY, FGFBP2, ZEB2, PRDM1, LAG3, FASLG, TRDC, CD38, MKI67* | *CD4, SLC4A10, IL6T, FOXO1, TCF7, CD27, IL7R, CD28, LEF1, FAS, CXCR4, CD44, IL2RA, KLF2, DPP4, CCR7, S1PR1, CD69, ABCB1, GATA3, BCL6, MAF, CXCR5, FOXP3, AHR, CCR6, RORC, IL23R, IL10RA, IL18R1, KLRG1, PDCD1* |

**Supplemental Table 7: T cell annotation marker genes (second level)**
